# Age-associated skeletal muscle dysfunction with spinal cord pathology in an *App* knock-in model of Alzheimer’s disease

**DOI:** 10.64898/2026.11.15.766852

**Authors:** Tsuyoshi Nakai, Yuya Urano, Akira Sobue, Mari Henmi, Shun Asai, Keisuke Hitachi, Taku Nagai, Hirohisa Watanabe, Koji Yamanaka, Kunihiro Tsuchida, Kiyofumi Yamada, Takashi Saito, Takaomi C. Saido, Shigeki Yamada, Atsushi Enomoto, Hiroyuki Mizoguchi

**Affiliations:** Department of Pharmacotherapeutics and Informatics, Fujita Health University School of Medicine, 1-98 Dengakugakubo, Kutsukake-cho, Toyoake, Aichi 470-1192, Japan; Department of Neuropsychopharmacology and Hospital Pharmacy, Nagoya University Graduate School of Medicine, 65 Tsurumai-cho, Showa-ku, Nagoya 466-8560, Japan; Department of Tumor Pathology, Nagoya University Graduate School of Medicine, 65 Tsurumai-cho, Showa-ku, Nagoya 466-8560, Japan; Division of Behavioral Neuropharmacology, International Center for Brain Science (ICBS), Fujita Health University, 1-98 Dengakugakubo, Kutsukake-cho, Toyoake, Aichi 470-1192, Japan; Fujita Mind-Brain Research & Innovation Center for Drug Generation (Fujita Mind-BRIDGe), Fujita Health University, 1-98 Dengakugakubo, Kutsukake-cho, Toyoake, Aichi 470-1192, Japan; Division for Therapies against Intractable Diseases, Center for Medical Science (CMS), Fujita Health University, 1-98 Dengakugakubo, Kutsukake-cho, Toyoake, Aichi 470-1192, Japan; Department of Neurology, Fujita Health University School of Medicine, 1-98 Dengakugakubo, Kutsukake-cho, Toyoake, Aichi 470-1192, Japan; International Center for Brain Science (ICBS), Fujita Health University, 1-98 Dengakugakubo, Kutsukake-cho, Toyoake, Aichi 470-1192, Japan; Department of Neuroscience and Pathobiology, Research Institute of Environmental Medicine, Nagoya University, Furo-cho, Chikusa-ku, Nagoya 464-8601, Japan; Institute for Glyco-Core Research (iGCORE), Nagoya University, Nagoya, Japan; Center for One Medicine Innovative Translational Research (COMIT), Nagoya University, Nagoya, Japan; Department of Neuropathology, Graduate School of Medicine, The University of Tokyo, 7-3-1 Hongo, Bunkyo-ku, Tokyo 113-0033, Japan; Laboratory for Proteolytic Neuroscience, RIKEN Center for Brain Science, 2-1 Hirosawa, Wako, Saitama 351-0198, Japan

**Keywords:** Alzheimer’s disease, Aging, Amyloid-β, Skeletal muscle, Spinal cord, *App*^NL-G-F/NL-G-F^ knock-in mice

## Abstract

Skeletal muscle dysfunction is increasingly recognized in Alzheimer’s disease (AD), but its pathological basis remains poorly understood. We examined age-associated muscle dysfunction together with skeletal muscle and spinal motor system alterations in male *App*^NL-G-F/NL-G-F^ knock-in (*App*-KI) mice, which develop age-dependent amyloid-β (Aβ) pathology without amyloid precursor protein (APP) overexpression. Middle-aged, but not adult, *App*-KI mice showed lower strength-related performance than age-matched wild-type mice, whereas skeletal muscle mass and myofiber cross-sectional area were preserved. General motor performance was also preserved, suggesting that the strength-related impairment did not simply reflect generalized motor dysfunction. Middle-aged *App*-KI mice had higher percentages of centrally nucleated fibers in the gastrocnemius and quadriceps femoris muscles. Transcriptomic and targeted expression analyses further revealed alterations in molecular profiles related to muscle development and myosin, including reduced expression of factors involved in myoblast fusion and selected myosin proteins. Despite evident cerebral Aβ pathology, no apparent Aβ immunoreactivity was detected in skeletal muscle. In the lumbar ventral horn, *App*-KI mice had fewer choline acetyltransferase-positive motor neurons, greater Aβ deposition, and increased immunoreactivity for glial fibrillary acidic protein and ionized calcium-binding adapter molecule 1; serum interleukin-6 levels were also selectively elevated in middle-aged *App*-KI mice. Together, these findings indicate that age-associated reductions in strength-related performance in *App*-KI mice occur without overt muscle atrophy and are accompanied by histological and molecular abnormalities in skeletal muscle and pathological changes in the lumbar spinal cord. Thus, AD-related physical dysfunction may involve abnormalities in both peripheral skeletal muscle and the spinal motor system.

## INTRODUCTION

Alzheimer’s disease (AD) is a progressive neurodegenerative disease and the most common cause of dementia [1]. Aging, a major risk factor for AD [1], is also accompanied by declining skeletal muscle mass and strength [2]. Physical dysfunction, particularly reduced muscle strength and impaired mobility, is common in patients with AD, and reduced muscle strength has been associated with incident AD, mild cognitive impairment, and cognitive decline [3–6]. Importantly, patients with early or mild AD may have reduced muscle strength despite relatively preserved muscle mass [4]. This dissociation suggests that AD-associated muscle dysfunction may not be explained solely by loss of muscle mass, yet its pathological basis remains poorly understood.

Although AD is characterized primarily by cerebral pathology, including extracellular amyloid-β (Aβ) plaques and intracellular neurofibrillary tangles [7], experimental models also show skeletal muscle abnormalities. In 3xTg-AD mice, several studies have reported skeletal muscle Aβ accumulation, impaired muscle strength and function, histological abnormalities, and age-related muscle loss [8–10]. Muscle atrophy accompanied by increased myostatin expression has also been reported in APP/PS1 mice [11]. These observations indicate that skeletal muscle can be affected in the setting of AD-related pathology, although their characteristics and underlying mechanisms may differ among models.

Potential contributors to AD-associated muscle dysfunction may extend beyond intrinsic skeletal muscle abnormalities. In Tg2576 mice, reduced cholinergic innervation of skeletal muscle [12] and spinal motor neuron degeneration with motor deficits [13] have been reported. Spinal Aβ deposition and glial abnormalities have also been observed in 5xFAD mice [14], and Aβ plaques have been identified in the lumbar spinal cord of *App*^NL-G-F/NL-G-F^ knock-in (*App*-KI) mice [15]. Collectively, these findings raise the possibility that skeletal muscle dysfunction associated with AD-related pathology may involve both muscle alterations and changes in the spinal motor system (Fig. 1). However, these accompanying skeletal muscle and spinal cord changes remain poorly characterized.

**Figure 1.**
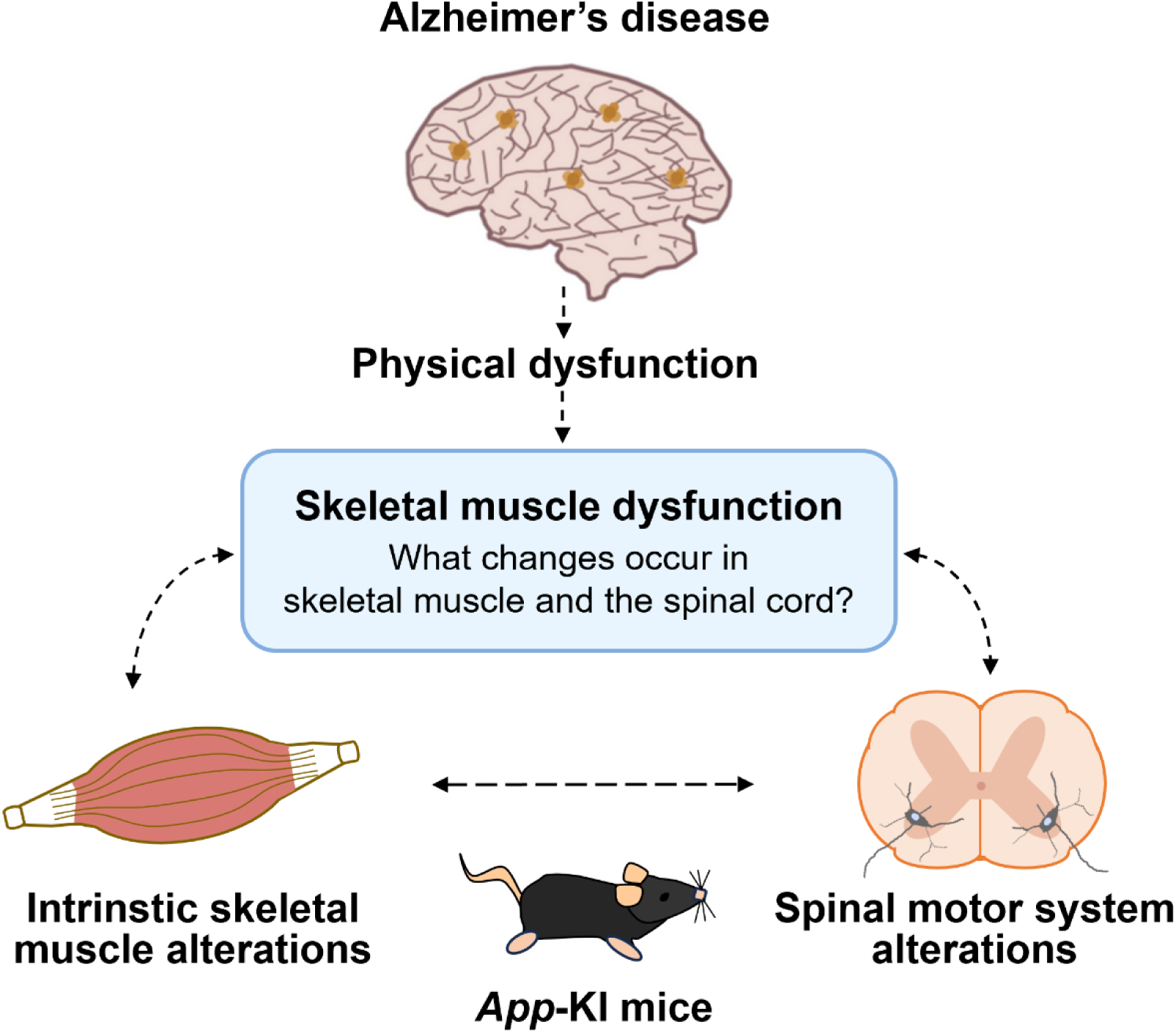
Conceptual diagram of skeletal muscle dysfunction in Alzheimer’s disease. Skeletal muscle dysfunction associated with Alzheimer’s disease may involve both intrinsic skeletal muscle alterations and changes in the spinal motor system. This working hypothesis guided the investigation of skeletal muscle and lumbar spinal cord phenotypes in *App* knock-in (*App*-KI) mice. Dashed arrows represent proposed relationships.

We therefore used *App*-KI mice to examine these changes in a model of Aβ pathology without amyloid precursor protein (APP) overexpression. These mice carry a humanized Aβ sequence together with the Swedish, Beyreuther/Iberian, and Arctic mutations in the endogenous *App* gene and develop age-dependent Aβ pathology, neuroinflammation and memory impairment [16]. The present study characterized age-associated skeletal muscle dysfunction in *App*-KI mice and examined the accompanying alterations in skeletal muscle and the lumbar spinal cord.

## MATERIALS AND METHODS

### Animals and tissues

*App*^NL-G-F^ knock-in mouse line (C57BL/6-*App*<tm3(NL-G-F)Tcs>) was originally generated as described by Saito et al. [16]. Homozygous *App*^NL-G-F/NL-G-F^ knock-in (*App*-KI) mice were obtained from the RIKEN BioResource Research Center (Tsukuba, Japan) and crossed with C57BL/6J mice purchased from CLEA Japan, Inc. (Tokyo, Japan) to generate heterozygous *App*^+/NL-G-F^ mice. These heterozygous mice were subsequently intercrossed to obtain homozygous *App*-KI mice and wild-type (WT) mice. The resulting *App*-KI and WT mice were maintained as separate breeding colonies, and age-matched WT mice were used as controls. *App*-KI and WT mice were housed under identical conditions (12-h light/dark cycle, 23 ± 3°C, and 50 ± 10 % relative humidity) with food and water available ad libitum. Animal experiments followed the guidelines established by the Japanese Pharmacological Society and the Institute for Laboratory Animals at Nagoya University and Fujita Health University. Protocols were approved by the Animal Care and Use Committee of Nagoya University (M210262-002) and Fujita Health University (APU25022-MD1). Only male mice were used to characterize age- and genotype-associated skeletal muscle and spinal cord phenotypes within a single sex. Adult *App*-KI mice aged 5–8 months and middle-aged *App*-KI mice aged 11–15 months were studied with age-matched WT controls; separate cohorts were used for the two age groups. Body weight was measured before each experiment, and skeletal muscle weights were recorded at tissue collection.

### Behavioral analyses

Behavioral assessments were conducted in WT and *App*-KI mice aged 5–6 months (adult) and 11–12 months (middle-aged). Adult mice underwent nest building, Y-maze, wire hanging, and grip strength tests, whereas middle-aged mice additionally underwent locomotor activity, rotarod, and treadmill exhaustion tests. To reduce bias from day-to-day variation in environmental conditions, WT and *App*-KI mice were examined in pairs on the same day. The experimenter was not blinded to the genotype during the behavioral assessments.

#### Nest building test

Nest building testing began between 9:00 and 10:00. Mice were housed individually in standard cages with clean sawdust and two 30.5 × 30.5 cm pieces of sterilized pulped virgin cotton-fiber nesting material (VWR International, LLC, Radnor, PA). Nest quality was assessed visually the following day using Deacon’s 5-point ordinal scale: 1 = nesting material not noticeably touched; 2 = nesting material partially torn; 3 = nesting material mostly shredded but without an identifiable nest site; 4 = an identifiable but flat nest; and 5 = a near-perfect or perfect nest [17].

### Y-maze test

The Y-maze test was performed as previously described [18], with minor modifications, under moderate illumination (35 lux) at the distal end of each arm. Each arm was 40 cm long, 12 cm high, 3 cm wide at the bottom, and 10 cm wide at the top. The arms converged to form an equilateral triangular central area measuring 4 cm along the longest axis. Each mouse was placed at the maze center and allowed to explore freely for 8 min while arm entries were recorded by visual observation. Spontaneous alternation was defined as consecutive entries into all three different arms within overlapping triplet sets. The alternation index (%) was calculated as [actual alternations/(total arm entries − 2)] × 100.

#### Grip strength test

Forelimb grip strength was measured as previously described [19], with minor modifications, using a grip strength meter (GST-M; O’Hara & Co., Tokyo, Japan). Each mouse grasped a wire grid with its forepaws and was gently pulled backward by the tail until release; peak forelimb force was recorded in newtons (N). Because the meter had an upper measurement limit of 1.0 N, values at or above this limit were recorded as 1.0 N. Three trials were performed at 30-min intervals, and the maximum of the three measurements was used for statistical analysis.

#### Wire hanging test

The wire hanging test was performed as previously described [19], with minor modifications, using a 22 × 22 × 33 cm box with a wire mesh ceiling (WH-3002; O’Hara & Co.). Each mouse was placed on the mesh before it was inverted, and latency to fall was recorded with a 60-s cutoff. Two trials were performed 1 h apart, and their mean was used for statistical analysis.

#### Locomotor activity test

Locomotor activity was assessed as previously described [18]. Each mouse was placed in a transparent rectangular rodent cage (25 × 30 × 18 cm) under moderate illumination (15 lux). Activity was recorded at 5-min intervals for 120 min using digital counters connected to infrared sensors positioned above the cage (NS-AS01; Brain Science Idea, Osaka, Japan).

#### Rotarod test

Rotarod testing was performed as previously described [20], with a shortened training period, and was conducted under moderate illumination (15 lux). Mice were trained for 3 consecutive days on a rod rotating at 6 rpm; latency to fall was recorded up to a maximum of 120 s. On day 4, mice were tested at 12 rpm. Six trials were conducted per day at 15-min intertrial intervals during both the training and test phases, and mean latency to fall was calculated for each mouse. The apparatus was cleaned with water and ethanol after every trial.

#### Treadmill exhaustion test

The treadmill exhaustion test was performed as previously described [21, 22], using a motorized single-lane treadmill (Columbus Instruments, Columbus, OH, USA) set at a 5° incline with a rear electrical stimulation grid delivering 0.5 mA. Mice were trained for 3 consecutive days at 5 m/min for 2 min, 7 m/min for 2 min, and 9 m/min for 1 min. After 1 day of rest, testing was performed on day 5, beginning at 5 m/min and increasing by 2 m/min every 2 min until exhaustion. Exhaustion was defined as remaining continuously on the rear stimulation grid for 10 s without resuming running. Total distance traveled before exhaustion was recorded for each mouse.

### Echocardiography

Transthoracic echocardiography was performed in 11–12-month-old (middle-aged) WT and *App*-KI mice as previously described [21]. Images were acquired with a Vevo 1100 system (FUJIFILM VisualSonics, Toronto, ON, Canada) equipped with an MS400 transducer (18–38 MHz). Parasternal short-axis and apical four-chamber views were obtained. Left ventricular (LV) end-systolic and end-diastolic volumes (LVESV and LVEDV) were measured from short-axis M-mode images at the mid-papillary muscle level, and peak early diastolic filling velocity (E) and peak early diastolic tissue velocity (e′) were also measured. Measurements were analyzed using Vevo LAB software (FUJIFILM VisualSonics). LV ejection fraction (EF) was calculated as EF (%) = (LVEDV − LVESV)/LVEDV × 100, and E/e′ was calculated by dividing E by e′.

### Aβ immunofluorescence staining in the brain

WT and *App*-KI mice aged 11–12 months (middle-aged) were deeply anesthetized with isoflurane and transcardially perfused with ice-cold phosphate-buffered saline (PBS), followed by 4% paraformaldehyde (PFA) in PBS. Brains were post-fixed, cryoprotected in 30% sucrose in PBS at 4°C until they sank, embedded in Tissue-Tek O.C.T. compound (#45833; Sakura Finetek,

Tokyo, Japan), and sagittally sectioned at 20 μm using an NX70 cryostat (Epredia Holdings Ltd., Portsmouth, NH, USA). Sections underwent antigen retrieval with HistoVT One (#06380-05; Nacalai Tesque, Kyoto, Japan) at 70°C for 20 min and were washed three times in PBS for 10 min each. After blocking for 1 h at room temperature (RT) with 5% normal donkey serum (#017-000-121; Jackson ImmunoResearch Laboratories, Inc., West Grove, PA, USA) in PBS containing 0.3% Triton X-100, sections were incubated overnight at 4°C with biotinylated mouse anti-Aβ N-terminal antibody (82E1; #10326; 1:200; Immuno-Biological Laboratories Co., Ltd.). Sections were then incubated for 1 h at RT with Alexa Fluor™ 633–conjugated streptavidin (1:500; Thermo Fisher Scientific, Waltham, MA, USA) and 4′,6-diamidino-2-phenylindole (DAPI; #D1306; 1:1,000; Thermo Fisher Scientific). Sections were mounted with Dako Fluorescence Mounting Medium (#S3023; Agilent Technologies), and cerebral cortical images were acquired using an LSM-980 confocal laser-scanning microscope with a 10× objective lens and ZEN software (Carl Zeiss AG, Oberkochen, Germany).

### Aβ immunofluorescence staining in skeletal muscle

Gastrocnemius (GC) and quadriceps femoris (QF) muscles from 11–12-month-old (middle-aged) WT and *App*-KI mice were oriented vertically on cork discs, secured with tragacanth gum, rapidly frozen in 2-methylbutane (#166-00615; FUJIFILM Wako Pure Chemical, Osaka, Japan) precooled with liquid nitrogen, and stored at −80°C. Muscles were cut into 10-μm transverse sections using a CM3000 cryostat (Leica Microsystems GmbH, Wetzlar, Germany). Sections were fixed in 4% PFA for 5 min at RT, washed three times in PBS, permeabilized with 0.3% Triton X-100 in PBS for 10 min, and blocked for 1 h with 5% normal donkey serum (Jackson ImmunoResearch Laboratories, Inc.) in PBS containing 0.3% Triton X-100. Sections were then incubated overnight at 4°C with the biotinylated mouse anti-Aβ N-terminal antibody (82E1; 1:200) described above. After washing, sections were incubated for 1 h at RT with Alexa Fluor™ 633–conjugated streptavidin (1:500; Thermo Fisher Scientific) and DAPI (1:1,000; Thermo Fisher Scientific), mounted with Dako Fluorescence Mounting Medium (Agilent Technologies), and imaged using an LSM-980 confocal laser-scanning microscope with a 10× objective lens and ZEN software (Carl Zeiss AG).

#### Laminin immunofluorescence staining and measurement of myofiber cross-sectional area

GC and QF muscles from WT and *App*-KI mice aged 6 months (adult) or 12 months (middle-aged) were mounted vertically on cork discs with tragacanth gum, rapidly frozen in 2-methylbutane (FUJIFILM Wako Pure Chemical) precooled with liquid nitrogen, and stored at −80°C. 10-μm-thick transverse sections were cut using a cryostat (Leica Microsystems GmbH), fixed in 4% PFA for 5 min at RT, and washed three times in PBS. Laminin immunostaining was performed using an Opal 3-Plex Manual Detection Kit (#NEL810001KT; Akoya Biosciences, Marlborough, MA, USA) according to the manufacturer’s instructions. After blocking with Antibody Diluent/Blocking (#ARD1001EA; Akoya Biosciences), sections were incubated for 1 h at RT with rabbit anti-laminin antibody (#L9393; 1:1,500; Sigma-Aldrich, St. Louis, MO, USA), washed four times with Tris-buffered saline with Tween 20 [TBS-T: 150 mM NaCl and 25 mM Tris–HCl (pH 7.5) with 0.05% Tween 20], and incubated for 10 min with Opal Polymer horseradish peroxidase secondary antibody (#ARH1001EA; Akoya Biosciences). After four further TBS-T washes, Opal 570 in 1 × Amplification Diluent (1:200) was applied for 10 min, followed by four washes and a 10-min incubation with Spectral DAPI in TBS-T (1:10) at RT. Sections were mounted using Epredia™ Immu-Mount™ (#9990412; Thermo-Fisher Scientific) and imaged using a BZ-X800 fluorescence microscope (Keyence, Osaka, Japan) with a 20× objective lens. Images within each staining batch were acquired using identical settings. An investigator blinded to genotype measured myofiber cross-sectional area with BZ-X Analyzer software (Keyence) in four randomly selected regions per muscle section; the mean across the four regions represented each mouse.

### Hematoxylin and eosin staining and quantification of centrally nucleated fibers

WT and *App*-KI mice aged 6 months (adult) or 12 months (middle-aged) were deeply anesthetized and transcardially perfused with ice-cold 0.1 M phosphate buffer (PB, pH 7.4), followed by 4% PFA in PB. GC and QF muscles were collected, paraffin-embedded, and cut into 4-μm transverse sections. Sections were deparaffinized, rehydrated through graded ethanol, stained with Gill’s hematoxylin, rinsed in water, counterstained with eosin, dehydrated, mounted, and coverslipped. Centrally nucleated fibers were quantified as previously described [23], with minor modifications. A nucleus was considered centrally located when it lay within the myofiber away from the sarcolemmal periphery. Images were acquired with a BZ-X800 microscope (Keyence) using a 20× objective lens. For each animal, 1,000 myofibers were evaluated per muscle type, and a fiber containing one or more centrally located nuclei was counted as centrally nucleated. The percentage of centrally nucleated fibers was calculated by dividing the number of centrally nucleated fibers by the total number of evaluated myofibers and multiplying by 100.

### Spinal cord immunofluorescence staining and image analysis

WT and *App*-KI mice aged 11–12 months (middle-aged) were deeply anesthetized with isoflurane and transcardially perfused with ice-cold PBS, followed by 4% PFA in PBS. Spinal cords were dissected, post-fixed in the same fixative overnight, cryoprotected in 30% sucrose in PBS at 4°C for 3 days, embedded in Tissue-Tek O.C.T. compound (Sakura Finetek), and stored at −80°C. Lumbar segments at approximately L3–L4 were cut into 20-μm transverse sections with a cryostat (Epredia Holdings Ltd.), and free-floating sections were used for staining. Separate sections underwent Aβ/choline acetyltransferase (ChAT) double immunostaining or Aβ/glial fibrillary acidic protein (GFAP)/ionized calcium-binding adaptor molecule 1 (IBA1) triple immunostaining. Sections were permeabilized with 0.5% Triton X-100 in PBS for 30 min and blocked for 1 h at RT in PBS containing 5% normal donkey serum (Jackson ImmunoResearch Laboratories, Inc.) and 0.3% Triton X-100. For Aβ/ChAT staining, sections were incubated overnight at 4°C with biotinylated mouse monoclonal anti-Aβ N-terminal antibody (82E1; 1:200) and goat polyclonal anti-ChAT antibody (#AB144P; 1:100; Sigma-Aldrich). For Aβ/GFAP/IBA1 staining, sections were incubated overnight at 4°C with 82E1 (1:200), rabbit anti-GFAP antibody (#MSFR101830; 1:250; Frontier Institute Co., Ltd., Hokkaido, Japan), and goat anti-AIF1/IBA1 antibody (#NB100-1028; 1:125; Novus Biologicals, Littleton, CO, USA). After three PBS washes, sections were incubated for 1 h at RT with Alexa Fluor™ 405–conjugated anti-goat IgG secondary antibodies (1:1,000; Thermo Fisher Scientific) and, for triple immunostaining, Alexa Fluor™ 594–conjugated anti-rabbit IgG secondary antibodies (1:1,000; Thermo Fisher Scientific), together with Alexa Fluor™ 633–conjugated streptavidin (1:500; Thermo Fisher Scientific). Sections were then washed and mounted with Dako Fluorescence Mounting Medium (Agilent Technologies). Images were acquired using an LSM-980 confocal laser-scanning microscope with a 20× objective lens and ZEN software (Carl Zeiss), using identical acquisition settings within each staining experiment. ImageJ software (National Institutes of Health, Bethesda, MD, USA) was used to quantify the percentages of Aβ-, GFAP-, and IBA1-immunopositive areas within predefined lumbar ventral horn regions of interest and to count ChAT-positive motor neurons. Six sections were analyzed per mouse for each parameter, and the mean across these sections represented as the value for each mouse. Image analysis was performed by an investigator blinded to genotype.

### RNA extraction and reverse transcription

Total RNA was extracted from GC muscles of 12-month-old (middle-aged) WT and *App*-KI mice using the RNeasy Mini Kit (#74104; Qiagen, Hilden, Germany), QIAGEN Proteinase K (#19131) and the RNase-Free DNase Set (#79254; Qiagen) according to the manufacturer’s instructions. Complementary DNA was synthesized from 0.5 μg total RNA using the ReverTra Ace qPCR RT Kit (#FSQ-101; TOYOBO, Osaka, Japan). Each 20-μL reverse transcription reaction contained 0.5 μg total RNA, 4 μL 5 × RT Master Mix, and nuclease-free water. Reverse transcription was performed at 37°C for 15 min, followed by enzyme inactivation at 98°C for 5 min using a MiniAmp Plus Thermal Cycler (Thermo Fisher Scientific).

### RNA sequencing and data analysis

RNA sequencing was performed by Rhelixa (Tokyo, Japan) using GC muscle from 12-month-old (middle-aged) WT and *App*-KI mice (*n* = 3 per group). Poly(A)-selected, strand-specific libraries were prepared from total RNA with the NEBNext^®^ Poly(A) mRNA Magnetic Isolation Module (#E7490; New England Biolabs, Ipswich, MA, USA) and NEBNext^®^ Ultra™ II Directional RNA Library Prep Kit (#E7760; New England Biolabs), then sequenced on an Illumina NovaSeq 6000 platform to generate 150-bp paired-end reads. Raw read quality was assessed with FastQC (version 0.11.7; https://www.bioinformatics.babraham.ac.uk/projects/fastqc/), and adapter sequences and low-quality bases were removed with Trimmomatic (version 0.38) [24]. Trimmed reads were aligned to the mouse reference genome (mm10) using HISAT2 (version 2.1.0) [25], and reads mapping to annotated exonic regions were quantified with featureCounts (version 1.6.3) [26]. Raw counts were normalized by the relative log expression method, and differential expression was analyzed with DESeq2 (version 1.24.0) [27]. For exploratory functional enrichment analysis, downregulated genes (fold change < 0.5, nominal *p* < 0.05) and upregulated genes (fold change > 2, nominal *p* < 0.05) were separately subjected to Gene Ontology Biological Process enrichment analysis using Metascape (https://metascape.org) [28]; terms with an enrichment factor > 1.5 and *p* < 0.01 were considered enriched. Heatmaps and MA plots were generated with the RIAS™ Visualization tool (Rhelixa).

### Quantitative real-time reverse transcription-PCR

Quantitative real-time reverse transcription-PCR was performed as previously described [21], with minor modifications. Each 20-μL reaction contained 10 μL THUNDERBIRD SYBR qPCR Master Mix (#QPS-201; TOYOBO), 1.2 μL each of 5 μM forward and reverse primers, 0.04 μL 50 × ROX reference dye, 7.06 μL nuclease-free water, and 0.5 μL cDNA template. Amplification was performed on an AriaMX Real-Time PCR System (Agilent Technologies Inc., Santa Clara, CA, USA) at 50°C for 120 s and 95 °C for 60 s, followed by 40 cycles of 95°C for 15 s and 60°C for 30 s. mRNA expression was calculated using the 2^−ΔΔ*Ct*^ method [29], normalized to β2-microglobulin (B2m), and expressed as fold change relative to WT controls. Primers were synthesized by Eurofins Genomics (Tokyo, Japan); sequences are listed in Supplementary Table 1.

### Western blotting

GC muscles from 12-month-old (middle-aged) WT and *App*-KI mice were sonicated in RIPA buffer (#182-02451; FUJIFILM Wako Pure Chemical) containing cOmplete™ protease inhibitor cocktail (#11873580001; Roche, Basel, Switzerland). Lysates were centrifuged at 15,000 × g for 20 min, and protein concentrations were measured with the DC Protein Assay Kit (#5000111JA; Bio-Rad Laboratories, Hercules, CA, USA). Samples were mixed with SDS-PAGE Sample Loading Buffer (#786-701; Geno Technology Inc., St. Louis, MO, USA) containing 5% 2-mercaptoethanol, heated at 95°C for 5 min, separated on 4–20% Mini-PROTEAN^®^ TGX™ Precast Protein Gels (#4561095; Bio-Rad), and transferred to polyvinylidene difluoride membranes (#IPVH00010; Merck Millipore, Billerica, MA, USA). Membranes were blocked with Blocking One-P (#05999-84; Nacalai Tesque) for 1 h at RT, incubated overnight at 4°C with primary antibodies, washed three times with TBS-T [10 mM Tris-HCl (pH 7.4), 100 mM NaCl, and 0.05% Tween-20], incubated for 1 h at RT with the appropriate HRP-conjugated secondary antibodies, and washed again three times. Immunoreactive bands were detected with ECL Prime Western Blotting Detection Reagent (#RPN2236; GE Healthcare, Chalfont St. Giles, Buckinghamshire, UK), imaged using a LuminoGraph I system (Atto Instruments, Tokyo, Japan), and quantified with CS Analyzer 4 software. Primary and secondary antibodies were diluted in Can Get Signal Solutions 1 and 2 (#NKB-101; Toyobo), respectively. Primary antibodies were mouse anti-MYH1 (#67299-1-Ig; 1:50,000; Proteintech Group, Rosemont, IL, USA), rabbit anti-MYH2 (#55069-1-AP; 1:600; Proteintech Group), rabbit anti-MYH4 (#20140-1-AP; 1:2,000; Proteintech Group), rabbit anti-MYH7 (#22280-1-AP; 1:500; Proteintech Group), rabbit anti-non-muscle myosin IIB/MYH10 (#8824T; 1:1,000; Cell Signaling Technology, Danvers, MA, USA), and rabbit anti-glyceraldehyde-3-phosphate dehydrogenase (GAPDH; #10494-1-AP; 1:2,500; Proteintech Group). HRP-conjugated anti-rabbit IgG (#7074; 1:1,000–10,000; Cell Signaling Technology) and anti-mouse IgG (#7076; 1:80,000; Cell Signaling Technology) served as secondary antibodies. GAPDH was the loading control; protein expression was normalized to GAPDH and expressed as fold change relative to WT controls.

### Measurement of serum creatine kinase and cytokines

Blood was collected from the inferior vena cava of deeply anesthetized mice aged 7–8 months (adult) or 14–15 months (middle-aged). Samples were allowed to clot for 30 min at RT, centrifuged at 2000 × g for 5 min, and the serum was aliquoted and stored at −80 °C. Serum creatine kinase (CK) activity was measured using FUJI DRI-CHEM slides CPK-PIII (#476-03301; FUJIFILM, Tokyo, Japan) and a DRI-CHEM 7000VZ automated analyzer (FUJIFILM). Serum interleukin-6 (IL-6) concentrations were measured with an enzyme-linked immunosorbent assay kit (#431301; BioLegend, San Diego, CA, USA). Serum IL-1β and tumor necrosis factor-α (TNF-α) were measured using a MILLIPLEX^®^ MAP Mouse Cytokine/Chemokine Magnetic Bead Panel (#MCYTOMAG-70K; Merck Millipore), a Luminex^®^200™ system, and xPONENT^®^ software (Luminex Corporation, Austin, TX, USA). All assays followed the manufacturers’ instructions.

### Statistical analysis

Except for RNA-sequencing data, statistical analyses were conducted in Prism 8.4.3 (GraphPad Software, La Jolla, CA, USA). Data are presented as mean ± standard error of the mean (SEM) unless otherwise stated. Normality and equality of variances were assessed using the Shapiro– Wilk and *F* tests, respectively. Two-group comparisons used an unpaired Student’s *t*-test for normally distributed data with equal variances, Welch’s *t*-test for normally distributed data with unequal variances, and the Mann–Whitney *U* test for non-normally distributed data. Nest building scores were compared between WT and *App*-KI mice within each age group using the Mann– Whitney *U* test. Y-maze spontaneous alternation and total arm entries, forelimb grip strength, wire hanging latency, body weight, relative skeletal muscle weights, and mean myofiber cross-sectional area were analyzed by two-way analysis of variance (ANOVA), with age and genotype as between-subject factors, followed by Bonferroni’s multiple comparisons test for WT versus *App*-KI comparisons within each age group. Locomotor activity and rotarod training performance were analyzed by two-way repeated-measures ANOVA, with genotype as the between-subject factor and time or training day as the within-subject factor; the Geisser–Greenhouse correction was applied. Total locomotor activity during the final 30 min and rotarod test-phase performance were analyzed using the Mann–Whitney *U* test, whereas treadmill performance was analyzed using an unpaired Student’s *t*-test. Serum CK activity and concentrations of IL-6, IL-1β, and TNF-α were compared between genotypes using the Mann–Whitney *U* test. All pairwise comparisons were two-sided, and *p* < 0.05 was considered statistically significant.

### AI-assisted manuscript preparation

ChatGPT (OpenAI, GPT-6) was used to assist English language editing and refinement of manuscript wording. Representative prompts requested improvements in clarity and naturalness while preserving the intended scientific meaning. All suggested revisions were reviewed by the authors before incorporation into the manuscript.

## RESULTS

### Age-associated behavioral and strength-related impairments in *App*-KI mice

Although *App*-KI mice develop age-dependent cerebral Aβ pathology, behavioral abnormalities have been reported at different ages and with varying severity across studies [16, 30, 31]. To determine whether behavioral abnormalities differed between adult and middle-aged *App*-KI mice, we first assessed nest building scores and Y-maze spontaneous alternation.

Nest building, an innate behavior sensitive to hippocampal and prefrontal cortical dysfunction [32, 33], is commonly used to assess functional impairments in AD mouse models [34–36]. Scores did not differ significantly between adult WT and *App*-KI mice (*p* = 0.0916) but were significantly lower in middle-aged *App*-KI mice than in age-matched WT mice (*p* < 0.0001; Fig. 2A). Y-maze spontaneous alternation, used here as a measure of spatial working memory [16, 37], showed a significant age × genotype interaction [*F*(1, 55) = 4.442, *p* = 0.0396] and a significant main effect of genotype [*F*(1, 55) = 19.47, *p* < 0.0001], with no significant main effect of age [*F*(1, 55) = 1.257, *p* = 0.2672]. Bonferroni testing showed no significant genotype difference in adults (*p* = 0.1994) but significantly lower alternation in middle-aged *App*-KI mice (*p* < 0.0001; Fig. 2B). Total arm entries showed a significant main effect of age [*F*(1, 55) = 15.69, *p* = 0.0002] but no significant main effect of genotype [*F*(1, 55) = 1.259, *p* = 0.2668] or age × genotype interaction [*F*(1, 55) = 1.022, *p* = 0.3165]; genotype comparisons were nonsignificant at both ages (adult, *p* = 0.2540; middle-aged, *p* > 0.9999; Fig. 2C).

**Figure 2.**
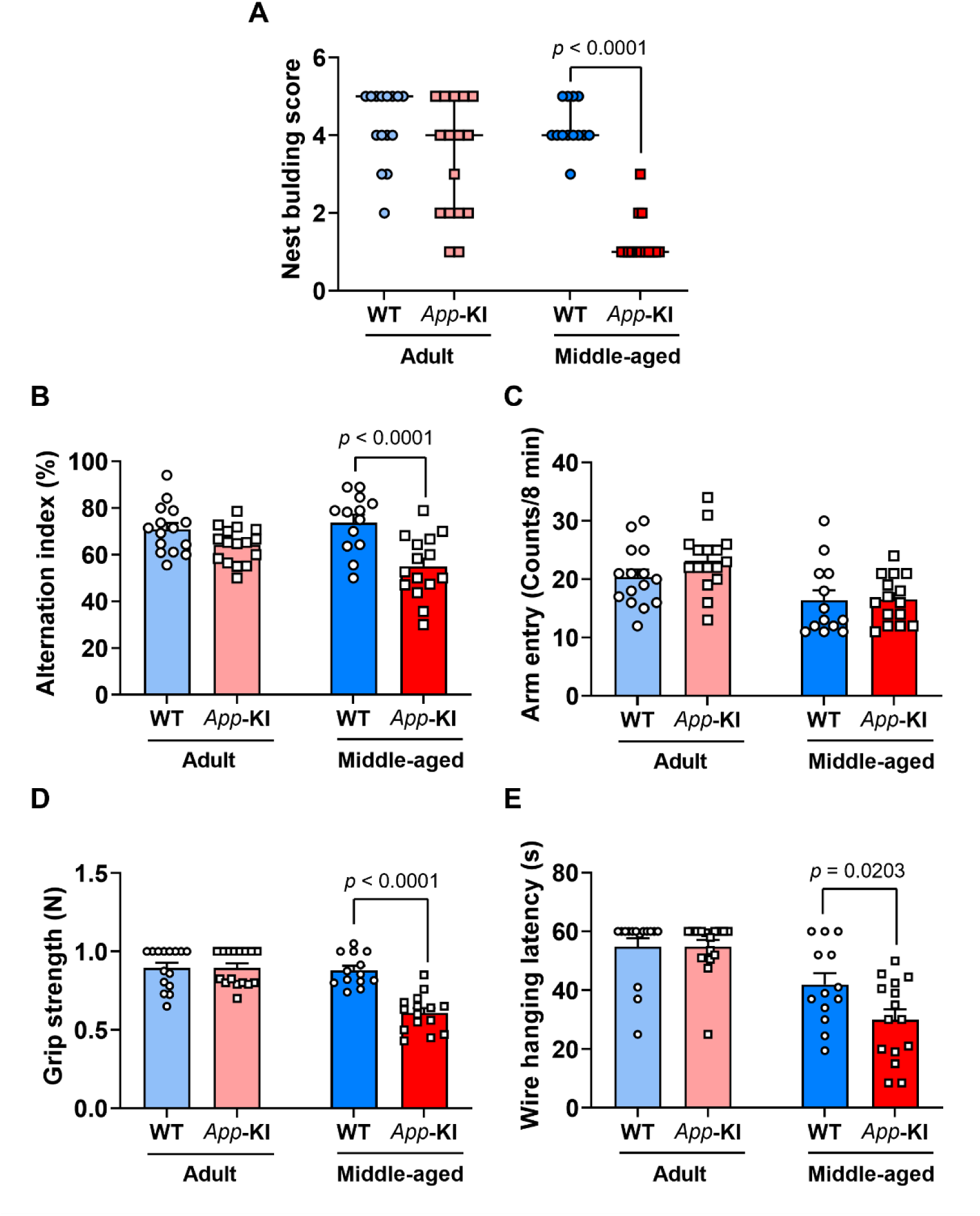
Age-associated behavioral and strength-related impairments in *App*-KI mice. (**A**) Nest building scores, (**B**) spontaneous alternation in the Y-maze test, (**C**) total arm entries in the Y-maze test, (**D**) forelimb grip strength, and (**E**) wire hanging latency in wild-type (WT) and *App*-KI mice aged 5–6 months (adult) and 11–12 months (middle-aged). Data in (A) are presented as the median with interquartile range, and data in (B–E) are expressed as the mean ± standard error of the mean (SEM); *n* = 15 (WT) and *n* = 16 (*App*-KI) for adult mice and *n* = 13 (WT) and *n* = 15 (*App*-KI) for middle-aged mice in (A–E). Nest building scores were analyzed using the Mann– Whitney *U* test within each age group. Data in (B–E) were analyzed using two-way analysis of variance (ANOVA) with age and genotype as between-subject factors, followed by Bonferroni’s multiple comparisons test to compare WT and *App*-KI mice within each age group.

We next evaluated strength-related performance using forelimb grip strength and wire hanging tests. Grip strength showed significant main effects of age [*F*(1, 55) = 25.27, *p* < 0.0001] and genotype [*F*(1, 55) = 20.33, *p* < 0.0001], together with a significant age × genotype interaction [*F*(1, 55) = 20.52, *p* < 0.0001]. Grip strength did not differ significantly between adult WT and *App*-KI mice (*p* > 0.9999) but was significantly lower in middle-aged *App*-KI mice than in age-matched WT mice (*p* < 0.0001; Fig. 2D). Wire hanging latency showed a significant main effect of age [*F*(1, 55) = 36.38, *p* < 0.0001], whereas the main effect of genotype [*F*(1, 55) = 3.723, *p* = 0.0588] and age × genotype interaction [*F*(1, 55) = 3.739, *p* = 0.0583] were not significant. Pairwise comparisons likewise showed no significant genotype difference in adults (*p* > 0.9999) but a significantly shorter latency in middle-aged *App*-KI mice (*p* = 0.0203; Fig. 2E). Thus, behavioral abnormalities and reduced strength-related performance emerged in middle-aged, but not adult, *App*-KI mice.

### Preserved general motor performance in middle-aged *App*-KI mice

To determine whether the strength-related deficits in middle-aged *App*-KI mice reflected a broader motor impairment, we assessed spontaneous locomotion, motor coordination and balance, and exercise endurance using locomotor activity, rotarod, and treadmill exhaustion tests, respectively [20–22].

Locomotor activity changed over time [*F*(5.773, 150.1) = 41.94, *p* < 0.0001], but neither the main effect of genotype [*F*(1, 26) = 1.041, *p* = 0.3169] nor the time × genotype interaction [*F*(5.773, 150.1) = 1.020, *p* = 0.4132] was significant (Supplementary Fig. 1A); total activity during the final 30 min also did not differ significantly between groups (*p* = 0.2788; Supplementary Fig. 1B). During rotarod training, latency to fall increased across days [*F*(1.814, 47.17) = 26.44, *p* < 0.0001] without a significant main effect of genotype [*F*(1, 26) = 0.0006613, *p* = 0.9797] or training day × genotype interaction [*F*(1.814, 47.17) = 1.492, *p* = 0.2357] (Supplementary Fig. 1C). Test-phase latency on day 4 did not differ significantly between groups (*p* = 0.2788; Supplementary Fig. 1D), and treadmill distance to exhaustion likewise did not differ significantly (*p* = 0.7390; Supplementary Fig. 1E). Because cardiac function can influence physical performance, we additionally examined echocardiographic parameters; neither EF nor E/e′ differed significantly between WT and *App*-KI mice (EF, *p* = 0.1611; E/e′, *p* = 0.8995; Supplementary Fig. 1F and G). These results indicate that no genotype differences were detected in the measured indices of general motor performance, endurance, or cardiac function in middle-aged *App*-KI mice. The behavioral and motor performance findings in adult and middle-aged mice are summarized in Table 1.

**Table 1.**
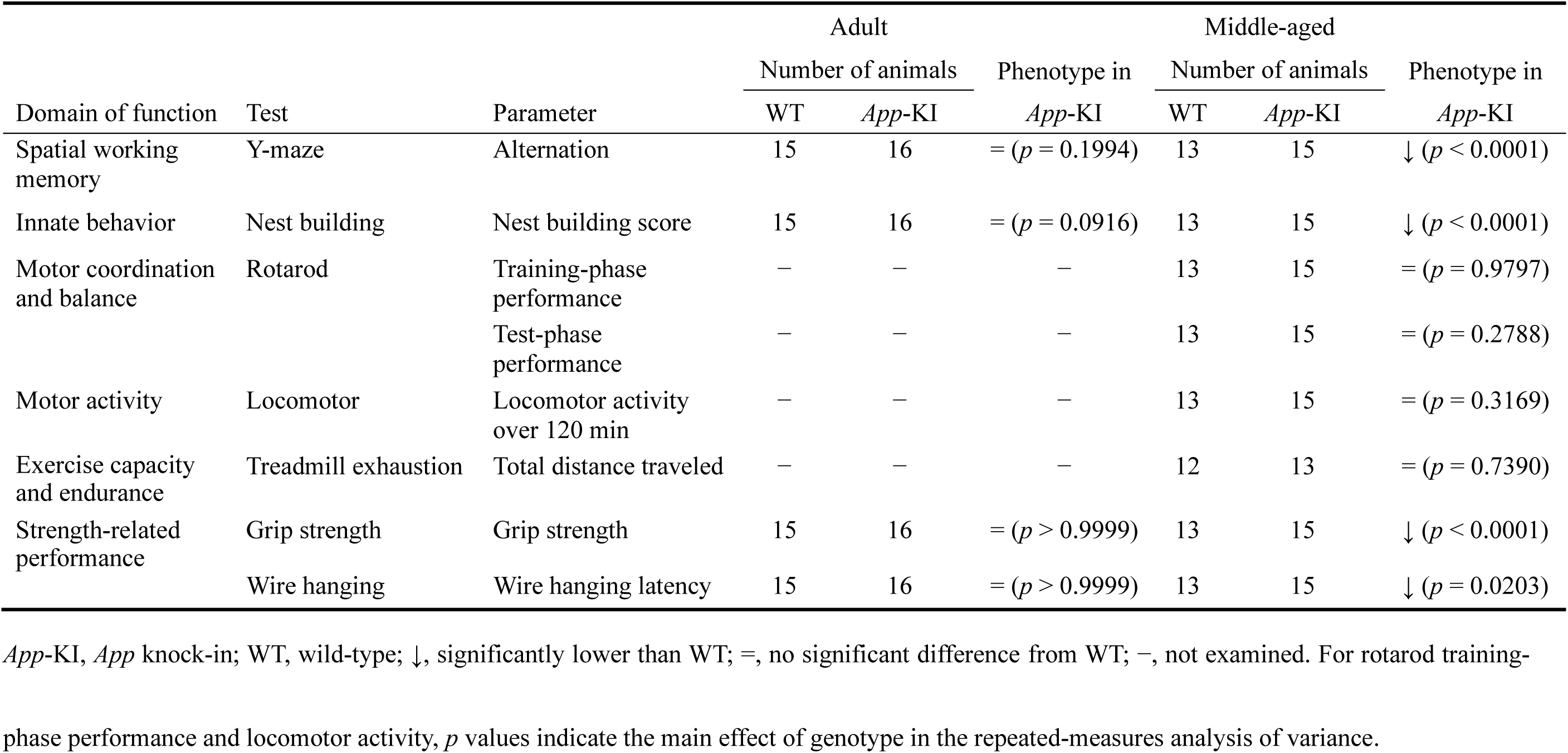
Summary of behavioral phenotypes in *App*-KI mice.

|  |  |  | Adult |  |  | Middle-aged |  |  |
|---|---|---|---|---|---|---|---|---|
|  |  |  | Number of animals |  | Phenotype in *App*-KI | Number of animals |  | Phenotype in *App*-KI |
| Domain of function | Test | Parameter | WT | *App*-KI |  | WT | *App*-KI |  |
| Spatial working memory | Y-maze | Alternation | 15 | 16 | = (*p* = 0.1994) | 13 | 15 | ↓ (*p* < 0.0001) |
| Innate behavior | Nest building | Nest building score | 15 | 16 | = (*p* = 0.0916) | 13 | 15 | ↓ (*p* < 0.0001) |
| Motor coordination and balance | Rotarod | Training-phase performance | − | − | − | 13 | 15 | = (*p* = 0.9797) |
|  |  | Test-phase performance | − | − | − | 13 | 15 | = (*p* = 0.2788) |
| Motor activity | Locomotor | Locomotor activity over 120 min | − | − | − | 13 | 15 | = (*p* = 0.3169) |
| Exercise capacity and endurance | Treadmill exhaustion | Total distance traveled | − | − | − | 12 | 13 | = (*p* = 0.7390) |
| Strength-related performance | Grip strength | Grip strength | 15 | 16 | = (*p* > 0.9999) | 13 | 15 | ↓ (*p* < 0.0001) |
|  | Wire hanging | Wire hanging latency | 15 | 16 | = (*p* > 0.9999) | 13 | 15 | ↓ (*p* = 0.0203) |

*App*-KI, *App* knock-in; WT, wild-type; ↓, significantly lower than WT; =, no significant difference from WT; −, not examined. For rotarod training-phase performance and locomotor activity, *p* values indicate the main effect of genotype in the repeated-measures analysis of variance.

### Preserved skeletal muscle mass and myofiber size in *App*-KI mice

We next asked whether reduced strength-related performance was accompanied by lower body weight, reduced muscle mass, or myofiber atrophy. Body weight showed a significant main effect of age [*F*(1, 36) = 18.65, *p* = 0.0001], but no significant main effect of genotype [*F*(1, 36) = 0.1856, *p* = 0.6691] or age × genotype interaction [*F*(1, 36) = 1.529, *p* = 0.2243] (Fig. 3A). Relative GC muscle weight showed no significant main effects of age [*F*(1, 36) = 2.960, *p* = 0.0939] or genotype [*F*(1, 36) = 0.1310, *p* = 0.7195] and no significant interaction [*F*(1, 36) = 0.7118, *p* = 0.4044] (Fig. 3B). Relative QF muscle weight showed a significant main effect of age [*F*(1, 36) = 16.12, *p* = 0.0003], but no significant main effect of genotype [*F*(1, 36) = 0.003197, *p* = 0.9552] or interaction [*F*(1, 36) = 0.3517, *p* = 0.5569] (Fig. 3C).

**Figure 3.**
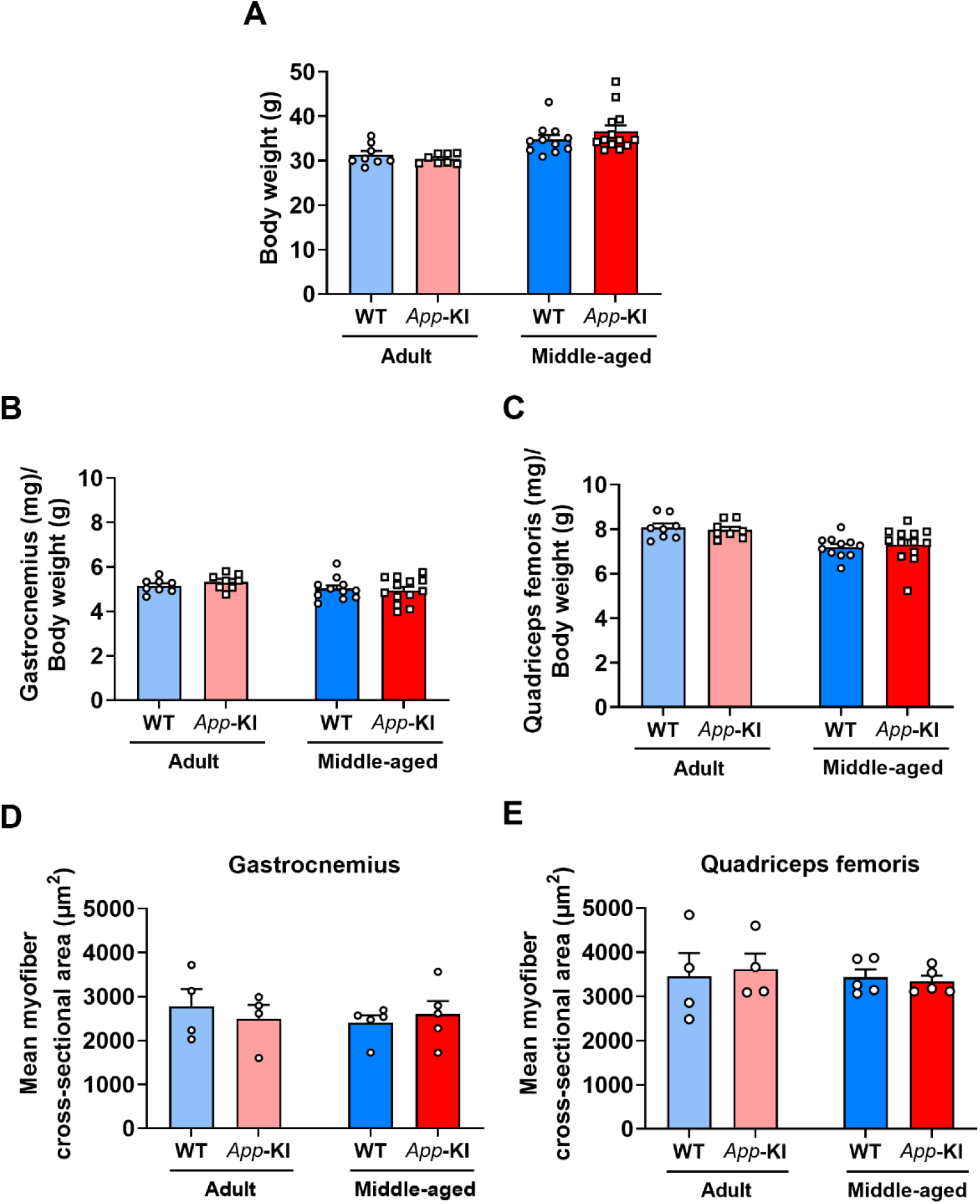
Preserved skeletal muscle mass and myofiber size in *App*-KI mice. (A) Body weight, (**B**) ratio of gastrocnemius (GC) muscle weight to body weight, (**C**) ratio of quadriceps femoris (QF) muscle weight to body weight, (**D**) mean myofiber cross-sectional area in the GC muscle, and (**E**) mean myofiber cross-sectional area in the QF muscle in wild-type (WT) and *App*-KI mice aged 6 months (adult) and 12 months (middle-aged). Data are presented as the mean ± SEM; *n* = 8 per group for adult mice and *n* = 11 (WT) and *n* = 13 (*App*-KI) for middle-aged mice in (A–C); *n* = 4 per group for adult mice and *n* = 5 per group for middle-aged mice in (D) and (E). Data were analyzed using two-way ANOVA with age and genotype as between-subject factors, followed by Bonferroni’s multiple comparisons test to compare WT and *App*-KI mice within each age group.

Mean myofiber cross-sectional area likewise showed no significant genotype effect. In GC, neither the main effect of age [*F*(1, 14) = 0.2295, *p* = 0.6393] nor the main effect of genotype [*F*(1, 14) = 0.01991, *p* = 0.8898] was significant, and no significant age × genotype interaction was detected [*F*(1, 14) = 0.6375, *p* = 0.4380] (Fig. 3D). The same pattern was observed in QF [age: *F*(1, 14) = 0.2445, *p* = 0.6287; genotype: *F*(1, 14) = 0.009789, *p* = 0.9226; interaction: *F*(1, 14) = 0.1793, *p* = 0.6784; Fig. 3E]. Bonferroni comparisons showed no significant genotype differences within either age group for body weight (adult, *p* > 0.9999; middle-aged, *p* = 0.3934), relative GC muscle weight (adult, *p* = 0.8818; middle-aged, *p* > 0.9999), relative QF muscle weight (both ages, *p* > 0.9999), or mean myofiber cross-sectional area (GC and QF at both ages, *p* > 0.9999). Myofiber size distributions largely overlapped between genotypes (Supplementary Figs. 2 and 3). Thus, no genotype-associated reductions in skeletal muscle mass or myofiber size were detected alongside the strength-related deficits in middle-aged *App*-KI mice.

### Histopathological alterations in skeletal muscle of middle-aged *App*-KI mice without apparent local Aβ immunoreactivity

Because the strength-related phenotype was not accompanied by overt muscle loss or myofiber atrophy (Figs. 2 and 3 and Supplementary Fig. 3), we next examined skeletal muscle histopathology. Centrally nucleated fibers, a feature commonly observed in regenerating myofibers after muscle injury, were quantified in the GC and QF muscles [38].

In adult mice, the percentage of centrally nucleated fibers did not differ significantly between WT and *App*-KI mice in either GC or QF (GC, *p* = 0.3802; QF, *p* = 0.1424; Supplementary Fig. 4). In middle-aged mice, however, this percentage was significantly higher in *App*-KI mice than in age-matched WT mice in both muscles (GC, *p* = 0.0001; QF, *p* < 0.0001; Fig. 4A–C). Serum CK, assessed as an additional indicator of muscle injury, was significantly higher in adult *App*-KI mice (*p* = 0.0260; Supplementary Fig. 5A) but did not differ significantly by genotype in middle-aged mice (*p* = 0.3527; Supplementary Fig. 5B).

**Figure 4.**
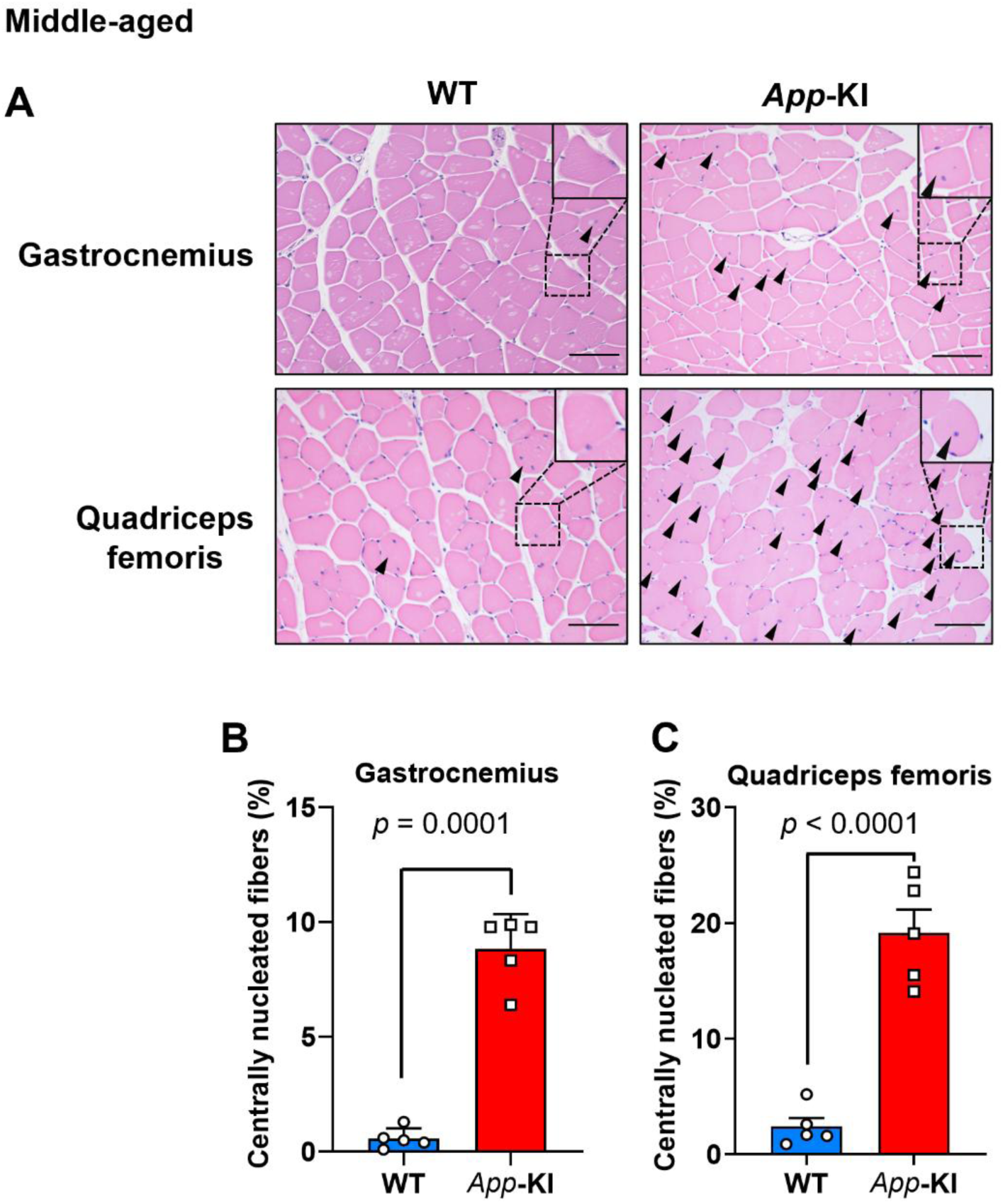
Increased percentages of centrally nucleated fibers in skeletal muscle from middle-aged *App*-KI mice. (**A**) Representative H&E-stained images of gastrocnemius (GC) and quadriceps femoris (QF) muscles from 12-month-old (middle-aged) wild-type (WT) and *App*-KI mice. Insets show higher-magnification views of the boxed regions. Arrowheads indicate centrally nucleated fibers. Scale bar: 100 μm. (**B, C**) Percentage of centrally nucleated fibers in the GC (B) and QF (C) muscles. Data are presented as the mean ± SEM; *n* = 5 per group, based on 1,000 myofibers per muscle per animal. Data were analyzed using Welch’s *t*-test for (B) and an unpaired Student’s *t*-test for (C). H&E, hematoxylin and eosin.

We then examined Aβ immunoreactivity in the brain and skeletal muscle of middle-aged mice. Cerebral cortex Aβ immunoreactivity was readily detected in *App*-KI mice but not in age-matched WT mice (Supplementary Fig. 6). By contrast, no apparent Aβ immunoreactivity was detected in the GC or QF muscles of *App*-KI mice. Thus, middle-aged *App*-KI mice displayed skeletal muscle histopathology despite preserved muscle mass and myofiber size and without apparent local Aβ immunoreactivity.

### Altered muscle-developmental- and myosin-related molecular profiles in middle-aged *App*-KI mice

To define the molecular changes accompanying this skeletal muscle phenotype, we performed genome-wide transcriptomic analysis of GC muscle from middle-aged WT and *App*-KI mice. Gene Ontology (GO) Biological Process enrichment analysis of genes downregulated in *App*-KI mice identified “muscle structure development” among the top 20 enriched terms (Fig. 5A). The corresponding analysis of upregulated genes is shown in Supplementary Fig. 7. Hierarchical clustering showed genotype-related differences in myosin- and myogenesis-related expression patterns (Fig. 5B). Several myosin-related genes, including *Myosin heavy chain 1* (*Myh1*), *Myh2*, *Myh3*, *Myh7*, and *Myh10*, showed lower expression in *App*-KI mice in the transcriptomic dataset (Fig. 5C); these genes encode fast- and slow-twitch, developmental, and non-muscle myosin isoforms [39, 40]. We therefore examined *myogenin* (*Myog*) and *myomaker* (*Mymk*), which contribute to myogenic differentiation and myoblast fusion, respectively [41, 42]. *Myog* expression did not differ significantly between WT and *App*-KI mice (*p* = 0.4355; Fig. 5D), whereas *Mymk* expression was significantly lower in *App*-KI GC muscle (*p* = 0.0440; Fig. 5E).

**Figure 5.**
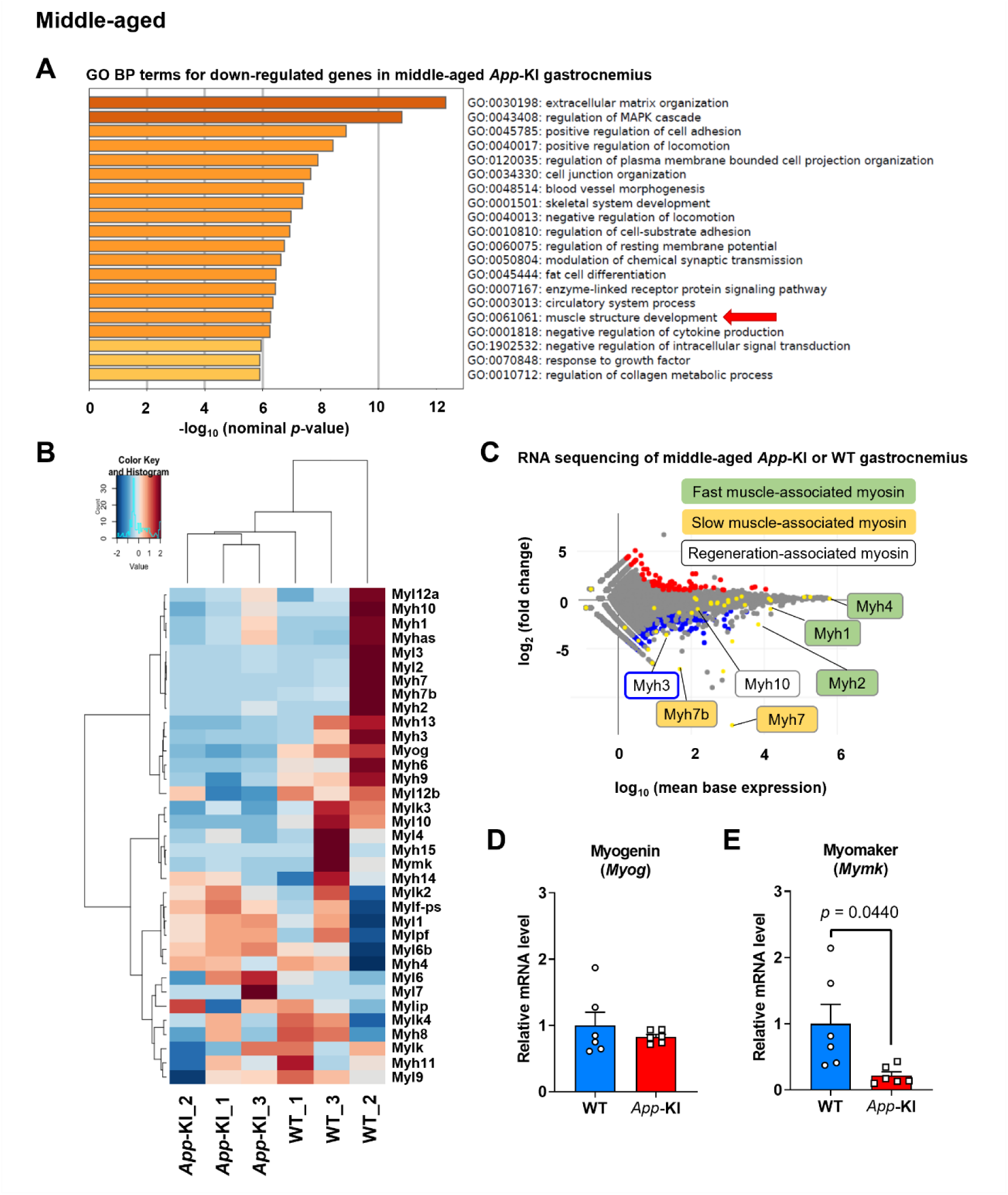
Altered muscle-development- and myosin-related gene expression in the gastrocnemius of middle-aged *App*-KI mice. (**A**) Gene Ontology (GO) biological process (BP) term enrichment analysis for downregulated genes in the gastrocnemius muscle of 12-month-old (middle-aged) *App*-KI mice. The top 20 terms are ranked by nominal *p* values. The red arrow highlights the “muscle structure development” term. (**B**) Heatmap and hierarchical clustering of myosin-encoding and myogenesis-related genes in 12-month-old (middle-aged) wild-type (WT) and *App*-KI mice (*n* = 3 per group). Color intensity represents each gene’s Z-score, as indicated in the key. (**C**) MA plot of gene expression changes in *App*-KI mice relative to WT mice. The y-axis represents the log2 fold change. Red and blue dots represent significantly upregulated and downregulated genes, respectively (adjusted *p* < 0.05). Yellow dots indicate myosin-encoding genes. Labeled myosin genes are classified as fast muscle-, slow muscle-, or regeneration-associated. (**D, E**) Relative expression of the muscle-regeneration-associated genes *myogenin* (*Myog*) (D) and *myomaker* (*Mymk*) (E) in gastrocnemius muscle of 12-month-old (middle-aged) WT and *App*-KI mice (*n* = 6 per group). Data in (D) and (E) are presented as the mean ± SEM and were analyzed using Welch’s *t*-tests.

We next tested whether the transcriptomic myosin changes were reflected at the protein level in middle-aged GC muscle; uncropped Western blots are shown in Supplementary Fig. 8. Among the fast-twitch-associated myosin isoforms examined, MYH1 protein expression did not differ significantly between genotypes (*p* = 0.1354; Fig. 6A), whereas MYH2 and MYH4 protein levels were significantly lower in *App*-KI mice (MYH2, *p* = 0.0331; MYH4, *p* = 0.0301; Fig. 6B and C). MYH7, a slow-twitch-associated isoform, did not differ significantly between genotypes (*p* = 0.3385; Fig. 6D), whereas MYH10, a non-muscle myosin IIB, showed significantly lower protein levels in *App*-KI mice (*p* = 0.0376; Fig. 6E). Overall, these findings suggest alterations in muscle-development- and myosin-related molecular profiles in middle-aged *App*-KI skeletal muscle, including reduced *Mymk* mRNA expression and lower levels of selected myosin proteins.

**Figure 6.**
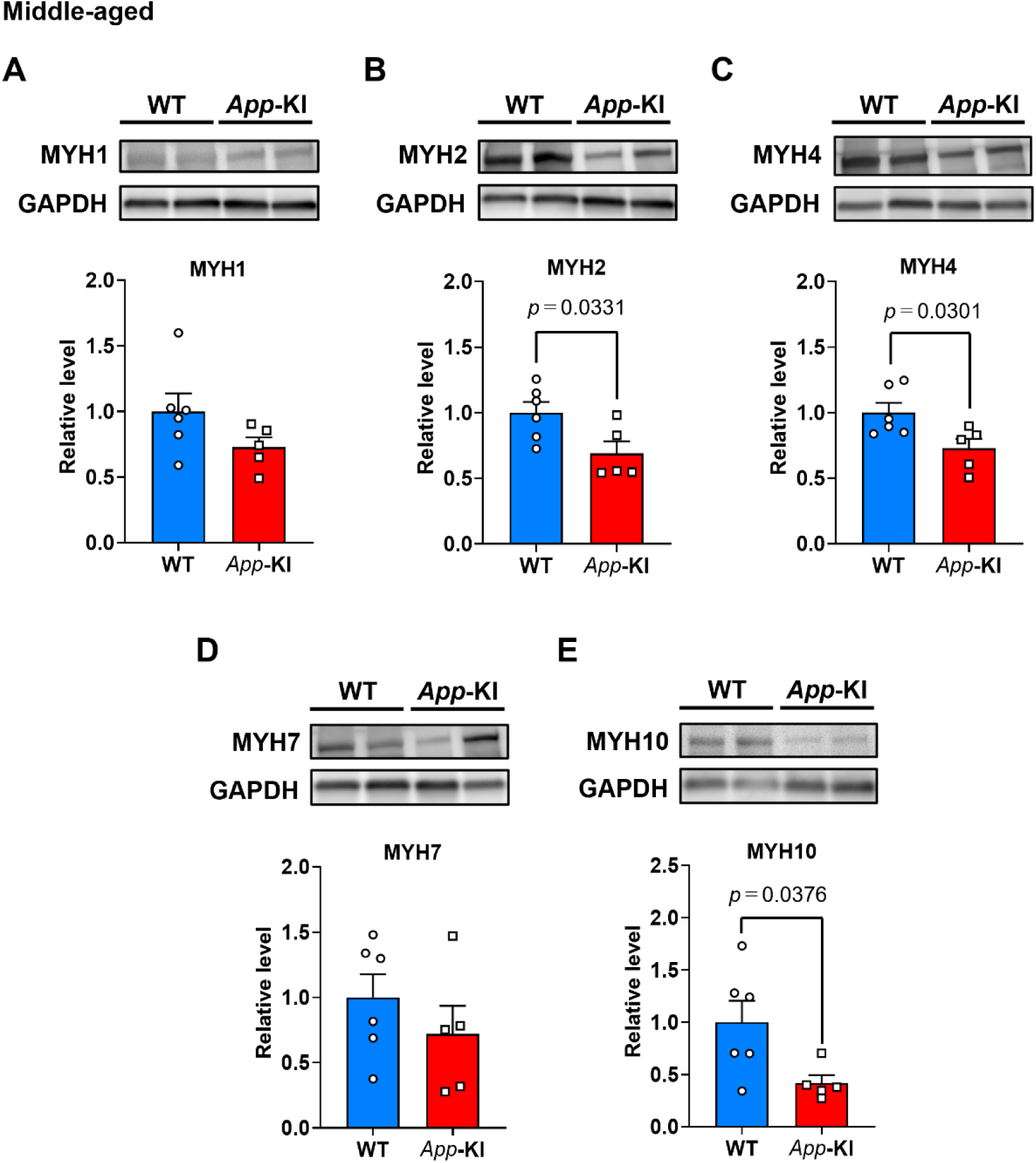
Altered myosin protein expression in the gastrocnemius muscle of middle-aged *App*-KI mice. (**A–E**) Representative immunoblots and quantification of MYH1 (A), MYH2 (B), MYH4 (C), MYH7 (D), and MYH10 (E) proteins in the gastrocnemius muscle of 12-month-old (middle-aged) wild-type (WT) and *App*-KI mice. Protein expression levels were normalized to GAPDH. Data are presented as the mean ± SEM; *n* = 6 (WT) and *n* = 5 (*App*-KI) for (A–E). Data were analyzed using unpaired Student’s *t*-tests. MYH1, myosin heavy chain 1; MYH2, myosin heavy chain 2; MYH4, myosin heavy chain 4; MYH7, myosin heavy chain 7; MYH10, non-muscle myosin heavy chain IIB; GAPDH, glyceraldehyde-3-phosphate dehydrogenase.

#### Fewer ChAT-positive motor neurons accompanied by Aβ deposition and increased GFAP and IBA1 immunoreactivity in the lumbar ventral horn of middle-aged *App*-KI mice

Because lumbar ventral horn motor neurons innervate hindlimb skeletal muscles, we examined whether the skeletal muscle phenotype was accompanied by lumbar spinal cord pathology. Middle-aged *App*-KI mice showed Aβ deposition in the ventral horn, with a significantly greater Aβ-immunopositive area than that in age-matched WT mice (*p* = 0.0058; Fig. 7A and B). The same region also contained fewer ChAT-positive motor neurons in *App*-KI mice (*p* = 0.0087; Fig. 7A and C).

**Figure 7.**
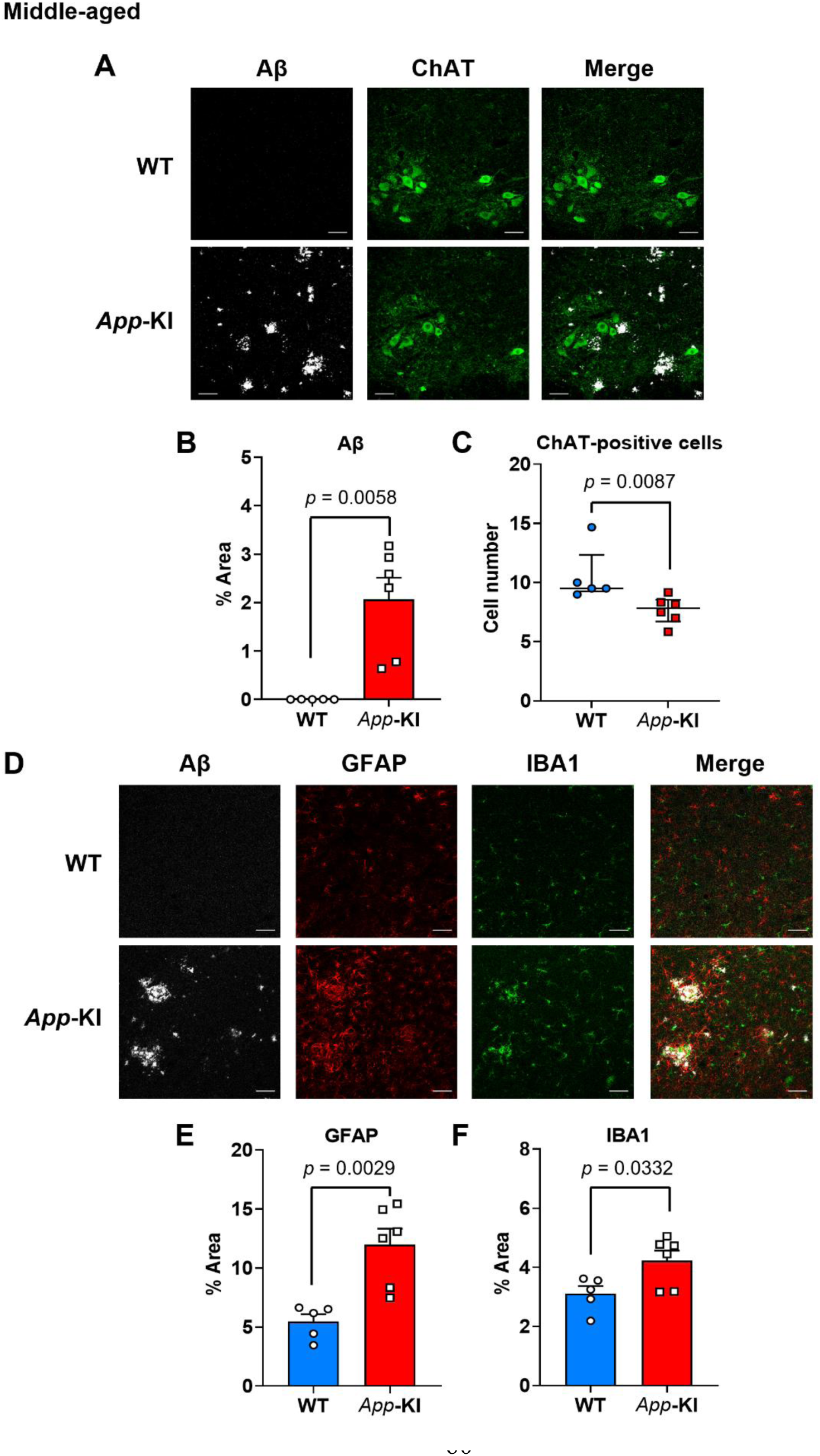
Fewer ChAT-positive motor neurons accompanied by Aβ deposition and increased GFAP and IBA1 immunoreactivity in the lumbar ventral horn of middle-aged *App*-KI mice. (**A**) Representative immunofluorescence images of amyloid-β (Aβ) and choline acetyltransferase (ChAT) in the lumbar ventral horn of 11–12-month-old (middle-aged) wild-type (WT) and *App*-KI mice. (**B, C**) Quantification of Aβ-immunopositive area (B) and ChAT-positive motor neurons (C) in the ventral horn. (**D**) Representative immunofluorescence images of Aβ, glial fibrillary acidic protein (GFAP), and ionized calcium-binding adaptor molecule 1 (IBA1) in the ventral horn. (**E, F**) Quantification of the GFAP-immunopositive area (E) and the IBA1-immunopositive area (F) in the ventral horn. Data are presented as the mean ± SEM for (B), (E), and (F), and as the median with interquartile range for (C); *n* = 5 (WT) and *n* = 6 (*App*-KI). For each parameter, six sections per mouse were analyzed, and the mean was used for statistical analysis. Scale bars: 50 μm. Data were analyzed using Welch’s *t*-test for (B), the Mann–Whitney *U* test for (C), and unpaired Student’s *t*-tests for (E) and (F).

We also assessed glial changes in the ventral horn by quantifying GFAP and IBA1 immunoreactivity as astrocytic and microglial markers, respectively. Both the GFAP-immunopositive area (*p* = 0.0029; Fig. 7D and E) and the IBA1-immunopositive area (*p* = 0.0332; Fig. 7D and F) were significantly greater in *App*-KI mice than in WT mice. Thus, the skeletal muscle abnormalities and reduced strength-related performance in middle-aged *App*-KI mice coincided with Aβ deposition, increased GFAP and IBA1 immunoreactivity, and fewer ChAT-positive motor neurons in the lumbar ventral horn.

### Circulating inflammatory changes in middle-aged *App*-KI mice

Because circulating proinflammatory cytokines have been implicated in AD-related inflammation [43, 44], we measured serum IL-6, IL-1β, and TNF-α. None differed significantly between 7–8-month-old (adult) WT and *App*-KI mice (IL-6, *p* = 0.3810; IL-1β, *p* = 0.4091; TNF-α, *p* = 0.9004; Supplementary Fig. 9A–C). In 14–15-month-old (middle-aged) mice, IL-6 was significantly higher in *App*-KI mice than in age-matched WT mice (*p* = 0.0002; Fig. 8A), whereas IL-1β and TNF-α did not differ significantly between genotypes (IL-1β, *p* = 0.0990; TNF-α, *p* = 0.6532; Fig. 8B and C). Thus, among the cytokines examined, only IL-6 was significantly elevated in middle-aged *App*-KI mice. The principal skeletal muscle, spinal cord, and circulating phenotypes are summarized in Table 2.

**Figure 8.**
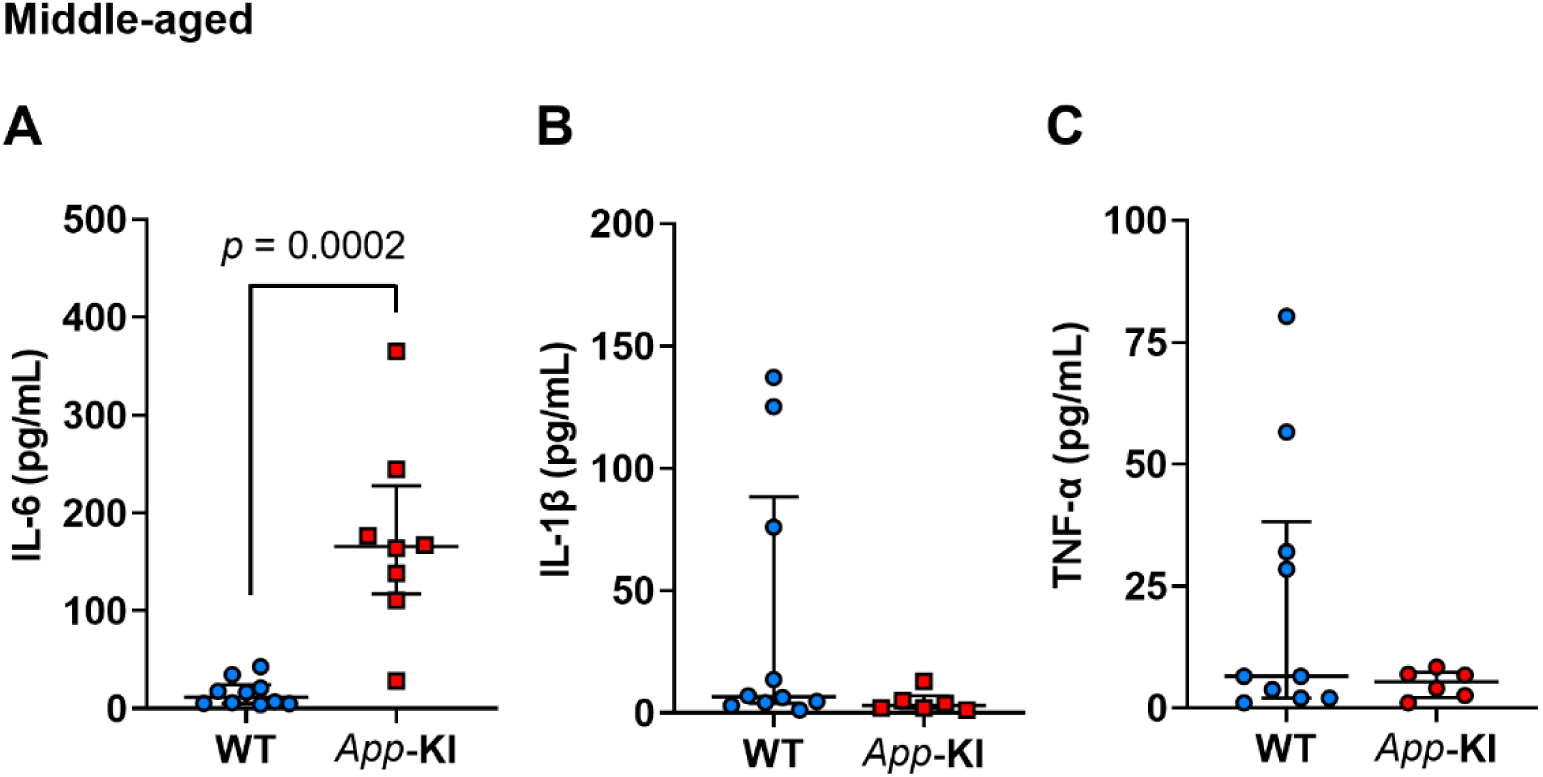
Increased serum IL-6 levels in middle-aged *App*-KI mice. Serum concentrations of. (**A**) IL-6, (**B**) IL-1β, and (**C**) TNF-α in 14–15-month-old (middle-aged) wild-type (WT) and *App*-KI mice. Data are shown as the median with interquartile range; *n* = 10 (WT) and *n* = 8 (*App*-KI) for (A), and *n* = 10 (WT) and *n* = 6 (*App*-KI) for (B) and (C). Data were analyzed using the Mann– Whitney *U* test. IL-6, interleukin-6; IL-1β, interleukin-1β; TNF-α, tumor necrosis factor-α.

**Table 2.** Summary of skeletal muscle, spinal cord, and circulating phenotypes in *App*-KI mice.

|  | Parameter | Phenotype in adult *App*-KI | Phenotype in middle-aged *App*-KI |
|---|---|---|---|
| Skeletal muscle | Relative skeletal muscle weight | = (GC, *p* = 0.8818; QF, *p* > 0.9999) | = (GC and QF, *p* > 0.9999) |
|  | Mean myofiber cross-sectional area | = (GC and QF, *p* > 0.9999) | = (GC and QF, *p* > 0.9999) |
|  | Centrally nucleated fibers | = (GC, *p* = 0.3802; QF, *p* = 0.1424) | ↑ (GC, *p* = 0.0001; QF, *p* < 0.0001) |
|  | mRNA expression |  |  |
|  | • *Mymk* | − | ↓ (*p* = 0.0440) |
|  | • *Myog* | − | = (*p* = 0.4355) |
|  | Protein expression |  |  |
|  | • MYH1 | − | = (*p* = 0.1354) |
|  | • MYH2 | − | ↓ (*p* = 0.0331) |
|  | • MYH4 | − | ↓ (*p* = 0.0301) |
|  | • MYH7 | − | = (*p* = 0.3385) |
|  | • MYH10 | − | ↓ (*p* = 0.0376) |
| Spinal cord | Immunofluorescence |  |  |
|  | • Aβ | − | ↑ (*p* = 0.0058) |
|  | • ChAT-positive motor neurons | − | ↓ (*p* = 0.0087) |
|  | • GFAP | − | ↑ (*p* = 0.0029) |
|  | • IBA1 | − | ↑ (*p* = 0.0332) |
| Serum | Circulating inflammatory markers |  |  |
|  | • IL-6 | = (*p* = 0.3810) | ↑ (*p* = 0.0002) |
|  | • IL-1β | = (*p* = 0.4091) | = (*p* = 0.0990) |
|  | • TNF-α | = (*p* = 0.9004) | = (*p* = 0.6532) |

*App*-KI, *App* knock-in; GC, gastrocnemius muscle; QF, quadriceps femoris muscle; ↑, significantly higher than wild-type (WT); ↓, significantly lower than WT; =, no significant difference from WT; −, not examined; *Mymk*, myomaker; *Myog*, myogenin; MYH1, myosin heavy chain 1; MYH2, myosin heavy chain 2; MYH4, myosin heavy chain 4; MYH7, myosin heavy chain 7; MYH10, non-muscle myosin heavy chain IIB; Aβ, amyloid-β; ChAT, choline acetyltransferase; GFAP, glial fibrillary acidic protein; IBA1, ionized calcium-binding adaptor molecule 1; IL-6, interleukin-6; IL-1β, interleukin-1β; TNF-α, tumor necrosis factor-α.

## DISCUSSION

The present study identifies an age-associated reduction in strength-related performance in *App*-KI mice without detectable genotype-associated reductions in skeletal muscle mass and myofiber size. In middle-aged *App*-KI mice, this functional phenotype was accompanied by increased centrally nucleated fibers and altered muscle-development- and myosin-related molecular profiles, and no apparent skeletal muscle Aβ immunoreactivity despite clear cerebral Aβ pathology. The lumbar ventral horn concurrently showed fewer ChAT-positive motor neurons, Aβ deposition, and increased GFAP and IBA1 immunoreactivity. Together, these findings suggest that reduced strength-related performance associated with AD-related pathology may involve abnormalities in both skeletal muscle and the spinal motor system, even in the absence of overt muscle wasting.

Middle-aged *App*-KI mice showed reduced strength-related performance, whereas no significant genotype differences were detected in the measured indices of general motor performance, exercise endurance, cardiac function, skeletal muscle mass, or myofiber size. Nest building and spatial working memory were also impaired in middle-aged, but not adult, *App*-KI mice. This pattern is consistent with clinical observations linking muscle strength to cognitive function: Filardi et al. reported lower handgrip strength across the AD clinical continuum and an association between handgrip strength and cognitive performance [5].

The dissociation between reduced strength-related performance and preserved muscle mass is another notable feature of this phenotype. Muscle strength often declines more rapidly than muscle mass during aging [45], and a similar dissociation has been observed in patients with AD. Ogawa et al. reported reduced muscle strength without a corresponding reduction in muscle mass in patients with early or mild AD, whereas both muscle strength and mass were reduced in patients with moderate AD [4]. AD mouse models likewise differ in their skeletal muscle phenotypes; aged APP/PS1 mice develop muscle atrophy associated with memory impairment [11], whereas the *App*-KI mice showed reduced strength-related performance and AD-like behavioral abnormalities without detectable reductions in muscle mass or myofiber size. This pattern may therefore parallel the dissociation between muscle strength and muscle mass described in patients with early or mild AD.

Direct analyses of skeletal muscle tissue from patients with AD remain limited, although a recent postmortem study reported molecular alterations in skeletal muscle [46]. In the present study, middle-aged *App*-KI mice showed histological and molecular alterations in skeletal muscle, without detectable genotype-associated reductions in muscle mass and myofiber size. Centrally nucleated fibers were increased in both GC and QF muscles. Centrally located nuclei are commonly observed in regenerated fibers after injury but can persist long after the initiating injury and therefore do not necessarily indicate ongoing or defective regeneration [38]. Serum CK activity was elevated in *App*-KI mice at 7–8 months and may reflect subtle muscle injury. However, because centrally nucleated fibers were assessed at 6 months, the relationship between these biochemical and histological findings remains uncertain. Transcriptomic analysis further revealed alterations in muscle-development- and myosin-related gene expression. Among the myogenic factors examined, *Mymk* expression was reduced, whereas *Myog* expression remained unchanged. Myomaker, encoded by *Mymk*, is essential for myoblast fusion and myofiber formation [41], whereas myogenin plays an important role in myogenic differentiation [42]. The selective reduction in *Mymk* expression may therefore reflect altered regulation of myoblast fusion-related processes. Together, these findings suggest that the skeletal muscle phenotype in middle-aged *App*-KI mice involves histological and molecular alterations rather than overt muscle atrophy.

Selective changes in myosin expression may also be relevant to this phenotype. Myosin heavy-chain isoforms are differentially expressed among skeletal muscle fiber types, with MYH1, MYH2, and MYH4 predominantly associated with fast-twitch fibers and MYH7 with slow-twitch fibers [39, 40]. *App*-KI mice showed significantly lower MYH2 and MYH4 protein levels, whereas MYH1 and MYH7 did not differ significantly between genotypes; MYH10, a non-muscle myosin IIB, was also significantly lower. The selective nature of these changes suggests alterations in specific myosin-related molecular components rather than a uniform reduction in myosin expression. These myosin-related changes may be related to reduced strength-related performance despite preserved muscle mass and myofiber size.

The absence of apparent local Aβ immunoreactivity distinguishes the *App*-KI muscle phenotype from several amyloidogenic models. Skeletal muscle Aβ accumulation has been reported in inclusion-body myositis and in experimental models expressing amyloidogenic APP- derived proteins [47, 48]. Increased skeletal muscle Aβ levels have also been reported in 3xTg-AD mice [8, 10], and reduced GC muscle mass has been observed in aged 3xTg-AD mice [10]. In contrast, no apparent Aβ immunoreactivity was detected in GC or QF muscles of *App*-KI mice despite the presence of histological and molecular muscle abnormalities. These findings suggest that local Aβ accumulation alone may not fully explain the muscle abnormalities observed in *App*-KI mice and that factors outside skeletal muscle may also contribute.

The concurrent lumbar spinal cord abnormalities represent one potential extramuscular component of the phenotype. Although human spinal cord pathology in AD is less well characterized than cerebral pathology [49], a postmortem study reported Aβ deposition in the spinal cord in some patients with AD, and postmortem studies have reported inconsistent evidence of spinal Aβ deposition [50, 51]. These findings suggest that spinal Aβ deposition may not be a uniform feature of AD. In the present study, the lumbar ventral horn in *App*-KI mice contained fewer ChAT-positive motor neurons together with Aβ deposition and increased GFAP and IBA1 immunoreactivity. Previous work in the same model reported spinal Aβ pathology and qualitative glial changes [15]; the present study extends those observations by quantifying GFAP and IBA1 immunoreactivity and ChAT-positive motor neurons while evaluating strength-related performance and skeletal muscle pathology in parallel. Other AD models also show heterogeneous spinal phenotypes: Tg2576 mice showed a reduction in lumbar spinal cord cholinergic neurons accompanied by motor abnormalities [13], whereas 5xFAD mice showed substantial spinal Aβ deposition and glial abnormalities, along with a reduction in cervical spinal cord neuronal nuclei-positive cells [14]. These observations suggest that spinal motor system alterations differ among AD mouse models. Although the present findings do not establish a causal relationship between spinal cord pathology and skeletal muscle dysfunction, their co-occurrence suggests that the spinal motor system may be relevant to at least some aspects of the strength-related phenotype observed in *App*-KI mice. Whether comparable spinal alterations occur in human AD and contribute to physical dysfunction remains unresolved.

A systemic inflammatory component may accompany these neural and muscle changes. Of the three circulating cytokines measured, only IL-6 was significantly elevated in middle-aged *App*-KI mice; IL-1β and TNF-α did not differ significantly between genotypes. Increased peripheral IL-6 levels have also been reported in patients with AD [44], and IL-6 has been suggested as a potential mediator of interactions between skeletal muscle and the central nervous system [43]. Sustained elevation of circulating IL-6 has also been reported to increase skeletal muscle fatigability without reducing muscle mass in mice [52]. The selective increase in circulating IL-6 is therefore consistent with a systemic inflammatory change accompanying the skeletal muscle and spinal cord abnormalities in middle-aged *App*-KI mice, although the functional relationship between these changes remains unclear.

This study has several limitations. First, the comparisons between adult and middle-aged mice were cross-sectional, so the temporal sequence of the behavioral, skeletal muscle, and spinal cord changes could not be established. Second, the concurrent lumbar spinal cord and muscle abnormalities do not demonstrate a causal relationship or define their relative contributions to reduced strength-related performance. Direct measurements of skeletal muscle contractile function and further evaluation of peripheral motor pathways could help clarify the mechanisms underlying the functional phenotype. Third, forelimb grip strength was assessed, whereas the histological analyses focused mainly on hindlimb skeletal muscles and the lumbar spinal cord. Although wire hanging performance was also reduced, this does not fully resolve the anatomical mismatch. In addition, some adult mice had grip strength values recorded at the upper measurement limit of 1.0 N, which may have reduced sensitivity to genotype differences in this age group. Finally, only male mice were studied, and whether similar phenotypes occur in female *App*-KI mice remains to be determined. Longitudinal and mechanistic studies integrating skeletal muscle, spinal and peripheral motor systems, and systemic factors are needed to define how these components contribute to AD-associated physical dysfunction.

## Acknowledgments

This work was supported by the Japan Society for the Promotion of Science (JSPS) KAKENHI Grant Numbers JP22K17824 and JP26K14596 (TN), the Japan Agency for Medical Research and Development (AMED) Grant Number JP21wm0425014 (K. Yamanaka), the Japan Society for Dementia Prevention (TN), the Nakatomi Foundation (TN), the Nitto Foundation (TN), and the Takeda Science Foundation (AS). The authors acknowledge the Division for Medical Research Engineering, Nagoya University Graduate School of Medicine, for access to a BZ-X800 microscope (KEYENCE), and the Fujita Health University Open Facility Center for access to an LSM-980 confocal laser scanning microscope with ZEN software (Carl Zeiss), a BZ-X800 microscope (KEYENCE), and a Luminex 200 system (Merck Millipore). We also thank the Division for Research of Laboratory Animals and the staff of the Division of Laboratory Animals at Nagoya University Graduate School of Medicine and the Advanced Medical Research Center for Animal Models of Human Diseases at Fujita Health University for their support. The authors used ChatGPT (OpenAI, GPT-6) to assist with English language editing and manuscript refinement. All AI-assisted suggestions were reviewed by the authors. We thank Editage (www.editage.jp) for English language editing.

## Authors’ Contributions

Conceptualization: Tsuyoshi Nakai and Hiroyuki Mizoguchi. Study design: Tsuyoshi Nakai and Yuya Urano. Methodology: Tsuyoshi Nakai, Yuya Urano, and Akira Sobue. Investigation: Tsuyoshi Nakai, Yuya Urano, Akira Sobue, Mari Henmi, Shun Asai, and Keisuke Hitachi. Formal analysis and data curation: Tsuyoshi Nakai. Visualization: Tsuyoshi Nakai and Yuya Urano. Resources: Taku Nagai, Takashi Saito, Takaomi C. Saido, Atsushi Enomoto, and Hiroyuki Mizoguchi. Funding acquisition: Tsuyoshi Nakai, Akira Sobue, Koji Yamanaka, and Hiroyuki Mizoguchi. Project administration: Tsuyoshi Nakai, Shigeki Yamada, and Hiroyuki Mizoguchi. Supervision: Taku Nagai, Hirohisa Watanabe, Koji Yamanaka, Kunihiro Tsuchida, Kiyofumi Yamada, Shigeki Yamada, Atsushi Enomoto, and Hiroyuki Mizoguchi. Writing – original draft: Tsuyoshi Nakai. Writing – review & editing: All authors. All the authors have read and approved the final version of the manuscript.

## Data availability

The RNA-sequencing data generated in this study have been deposited in the NCBI Gene Expression Omnibus (GEO) under accession number GSE346165. Other data supporting the findings of this study are available within the article and its Supplementary Data or from the corresponding authors upon reasonable request.

