## Supplementary Material for "Age-associated skeletal muscle dysfunction with spinal cord pathology in an *App* knock-in model of Alzheimer’s disease"

SUPPLEMENTARY DATA

**Supplementary Table 1. Primer sequences used for real-time RT-PCR.**

| Gene | NCBI accession number | Forward primer sequence (5'–3') | Reverse primer sequence (5'–3') | Product size (bp) |
|---|---|---|---|---|
| *B2m* | NM_009735.3 | ATACGCCTGCAGAGTTAAGCA | TCACATGTCTCGATCCCAGTAG | 70 |
| *Mymk* | NM_025376.3 | CATGTTCTTTGTGGCGTTCTCCC | AGTACTCCAGAATGTCACGGC | 86 |
| *Myog* | NM_031189.2 | GTCCCAACCCAGGAGATCATT | GCTGTCCACGATGGACGTAA | 83 |

RT-PCR, reverse transcription polymerase chain reaction; NCBI, National Center for Biotechnology Information; *B2m*, β2-microglobulin; *Mymk*, myomaker, myoblast fusion factor; *Myog*, myogenin.

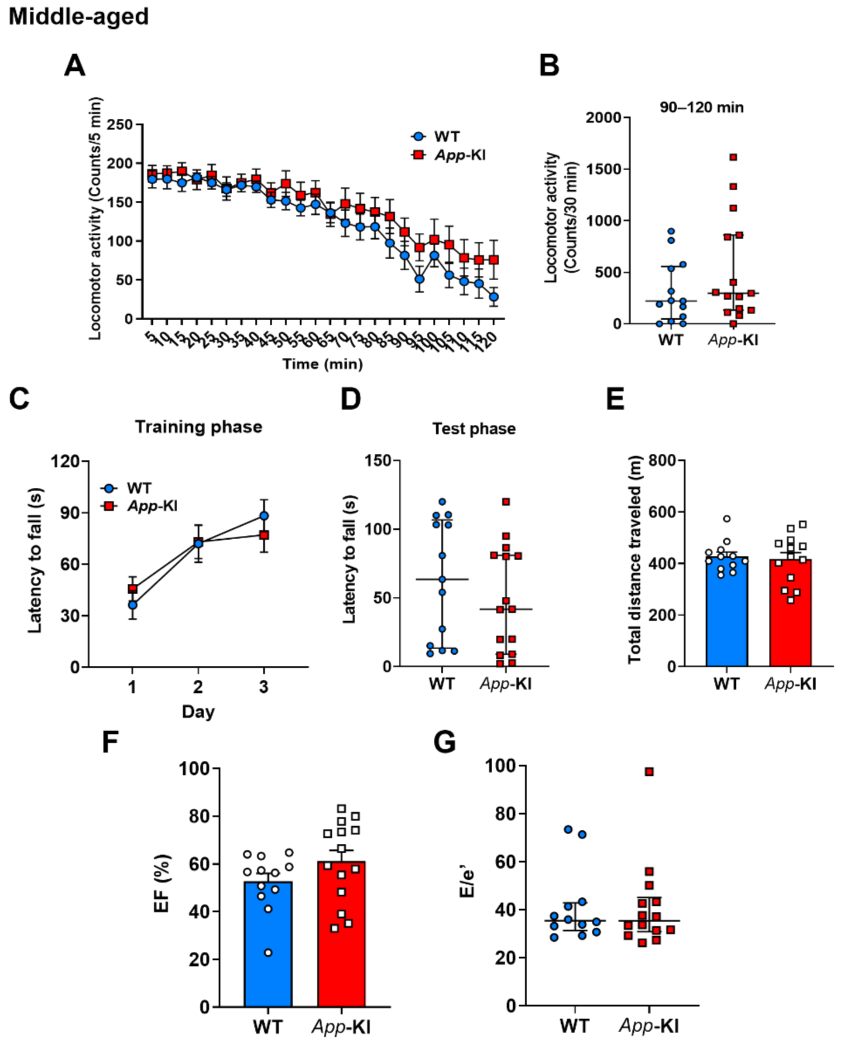


**Supplementary Figure 1. Preserved general motor performance and cardiac function in middle-aged *App*-KI mice.** General motor performance and cardiac function were assessed in wild-type (WT) and *App* knock-in (*App*-KI) mice aged 11–12 months (middle-aged). (**A**) Locomotor activity over 120 min. (**B**) Total locomotor activity during the final 30 min (90–120 min). (**C, D**) Rotarod performance during the training phase (days 1–3; C) and test phase (day 4; D). (**E**) Total distance traveled in the treadmill exhaustion test. (**F, G**) Echocardiographic assessment of ejection fraction (EF) (F) and the ratio of early mitral inflow velocity to early diastolic mitral annular velocity (E/e′) (G). Data are expressed as the mean ± standard error of the mean (SEM) for (A), (C), (E), and (F), and as median with interquartile range for (B), (D), and (G); $n = 13$ (WT) and $n = 15$ (*App*-KI) for (A–D), $n = 12$ (WT) and $n = 13$ (*App*-KI) for (E), and $n = 12$ (WT) and $n = 14$ (*App*-KI) for (F) and (G). Locomotor activity (A) and rotarod training performance (C) were analyzed using two-way repeated-measures analysis of variance with the Geisser–Greenhouse correction. Data in (B), (D), and (G) were analyzed using Mann–Whitney *U* tests, and (E) and (F) using unpaired Student's *t*-tests.

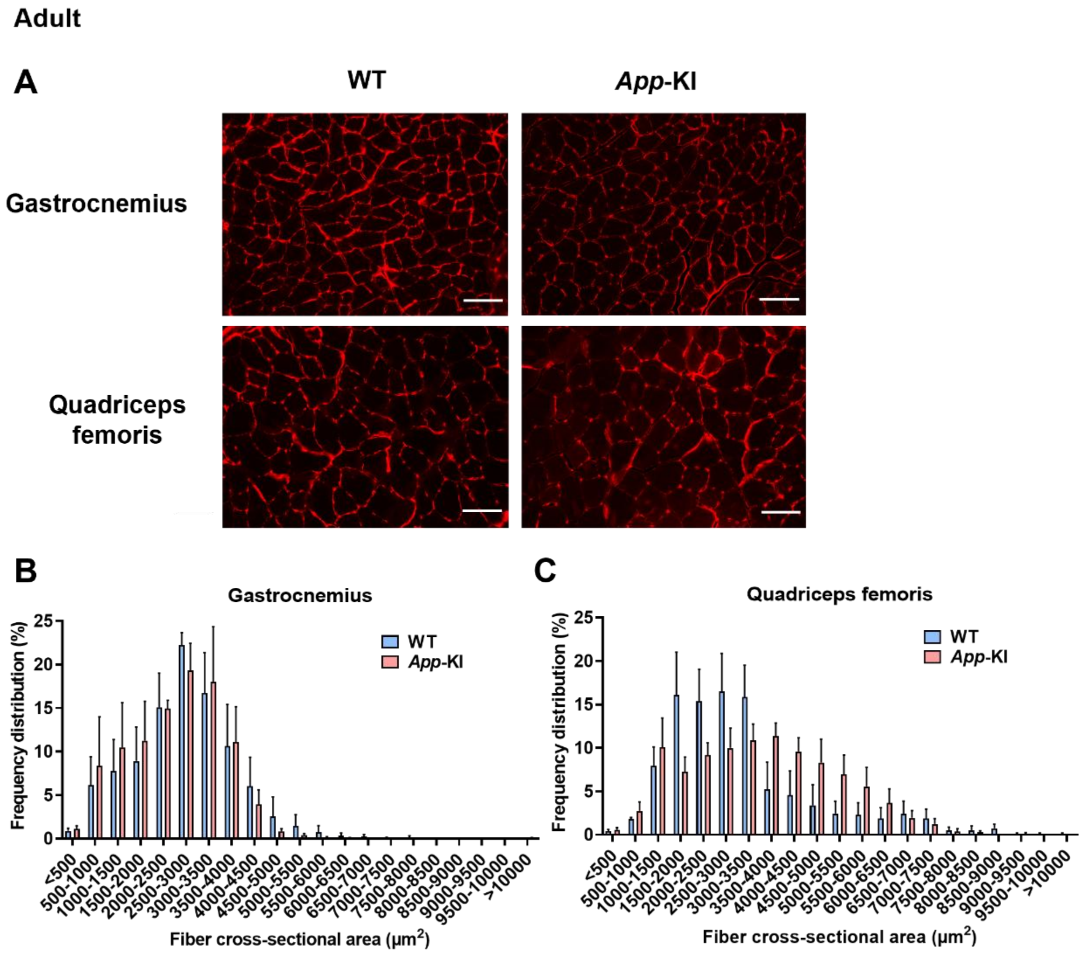


**Supplementary Figure 2. Myofiber size distribution in adult *App*-KI mice.** (**A**) Representative laminin-immunostained images of gastrocnemius (GC) and quadriceps femoris (QF) muscles from 6-month-old (adult) wild-type (WT) and *App*-KI mice. Scale bar: 100 µm. (**B, C**) Distribution of myofiber cross-sectional area in the GC (B) and QF (C) muscles. Data are presented as the mean ± SEM; *n* = 4 per group. Four randomly selected regions were analyzed per muscle section per animal.

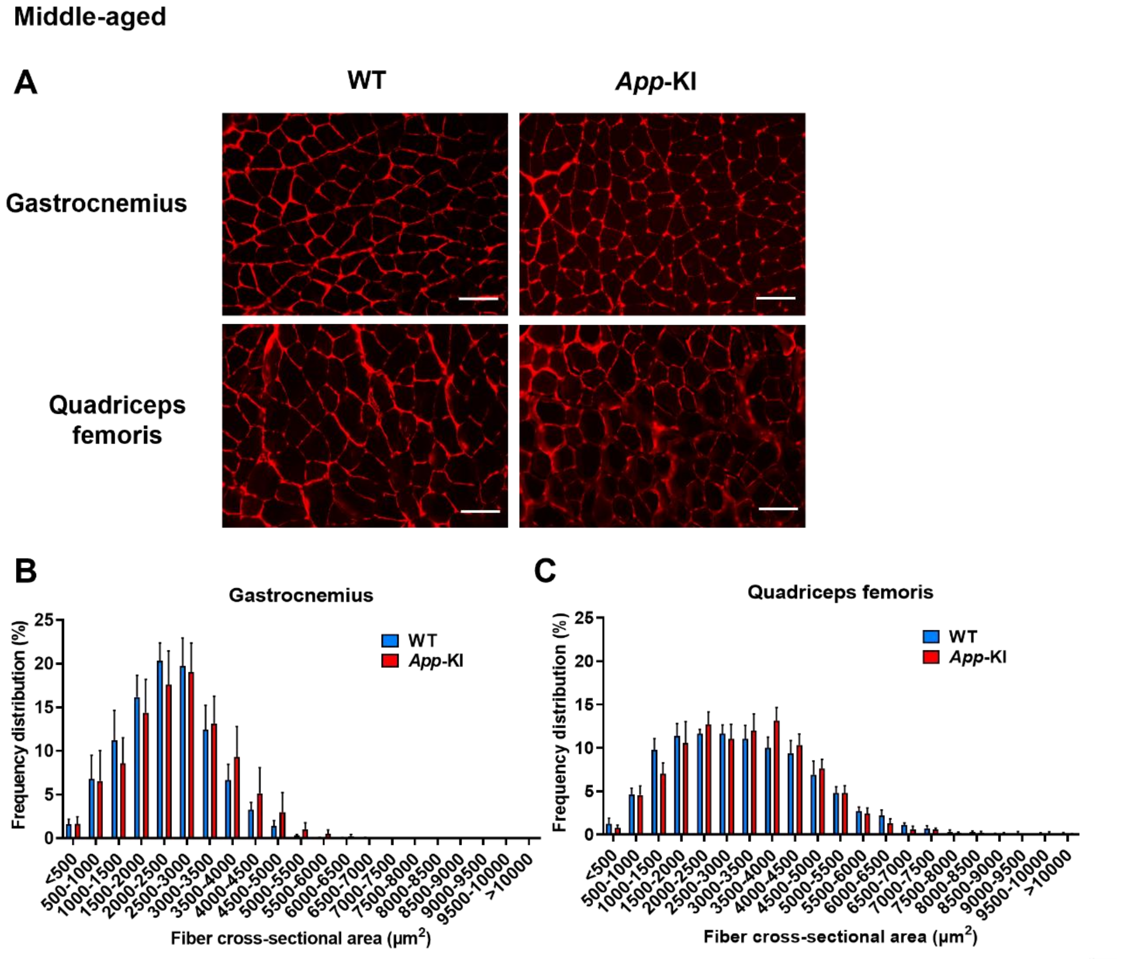


**Supplementary Figure 3. Myofiber size distribution in middle-aged *App*-KI mice.** (**A**) Representative laminin-immunostained images of gastrocnemius (GC) and quadriceps femoris (QF) muscles from 12-month-old (middle-aged) wild-type (WT) and *App*-KI mice. Scale bar: 100 µm. (**B, C**) Distribution of myofiber cross-sectional area in the GC (B) and QF (C) muscles. Data are presented as the mean ± SEM; *n* = 5 per group. Four randomly selected regions were analyzed per muscle section per animal.

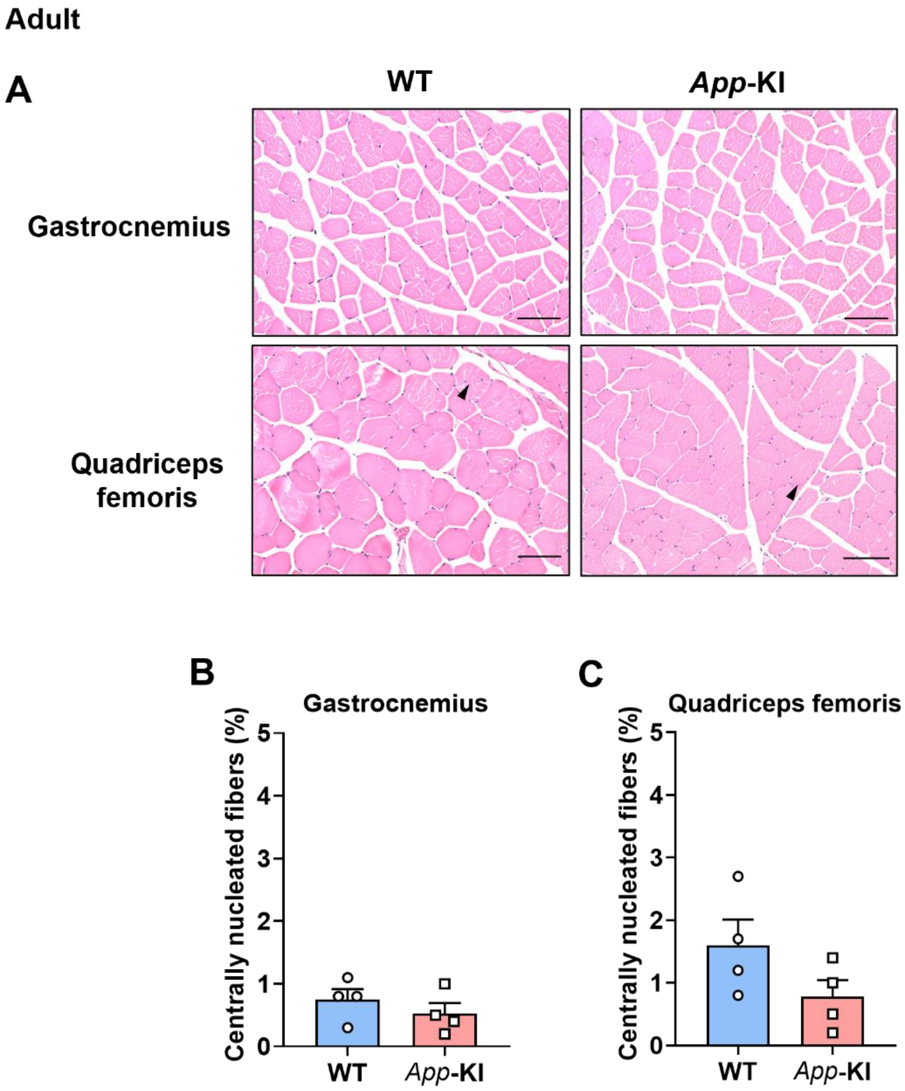


**Supplementary Figure 4. No increase in centrally nucleated fibers in skeletal muscle from adult *App*-KI mice.** (**A**) Representative H&E-stained images of gastrocnemius (GC) and quadriceps femoris (QF) muscles from 6-month-old (adult) wild-type (WT) and *App*-KI mice. Arrowheads indicate centrally nucleated fibers. Scale bar: 100 µm. (**B, C**) Percentage of centrally nucleated fibers in the GC (B) and QF (C) muscles. Data are presented as the mean ± SEM; $n = 4$ per group, based on 1,000 myofibers per muscle per animal. Data were analyzed using unpaired Student's $t$-tests. H&E, hematoxylin and eosin.

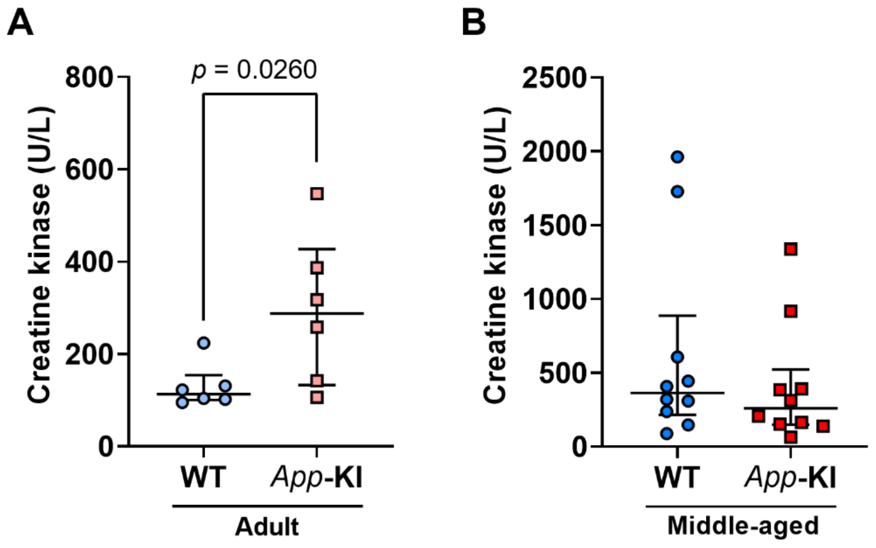


**Supplementary Figure 5. Serum creatine kinase activity in adult and middle-aged *App*-KI mice.** Serum creatine kinase (CK) activity in wild-type (WT) and *App*-KI mice aged 7–8 months (adult, panel **A**) and 14–15 months (middle-aged, panel **B**). Data are presented as the median with interquartile range; $n = 6$ per group in (A) and $n = 10$ per group in (B). Data were analyzed using the Mann–Whitney $U$ test.

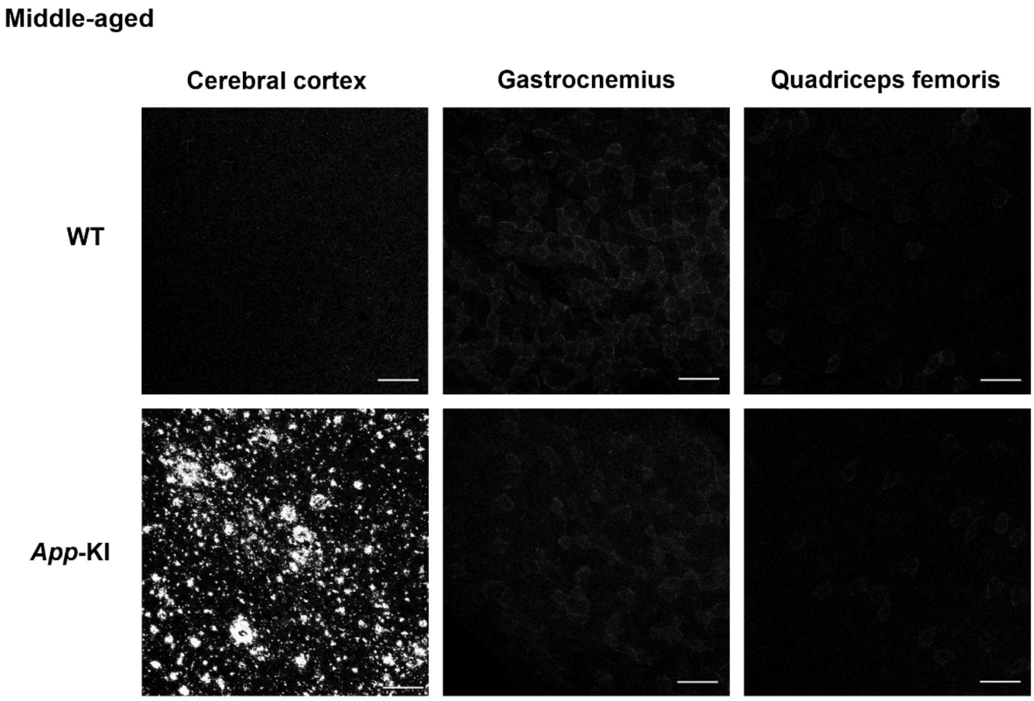


**Supplementary Figure 6. Aβ immunoreactivity in the brain and skeletal muscle of middle-aged wild-type (WT) and *App*-KI mice.** Representative immunofluorescence images of Aβ in the cerebral cortex, gastrocnemius, and quadriceps femoris muscles of 11–12-month-old (middle-aged) WT and *App*-KI mice. Scale bars: 100 µm. Aβ, amyloid-β.

### Middle-aged

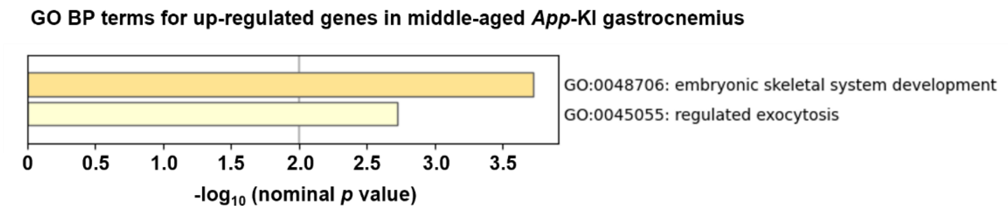


**Supplementary Figure 7. Gene Ontology biological process enrichment analysis of upregulated genes in the gastrocnemius of middle-aged *App*-KI mice.** Gene Ontology (GO) biological process (BP) term enrichment analysis for upregulated genes in the gastrocnemius muscle of 12-month-old (middle-aged) wild-type (WT) and *App*-KI mice ($n$ = 3 per group). Representative GO BP terms from the two clusters are ranked by nominal $p$ values.

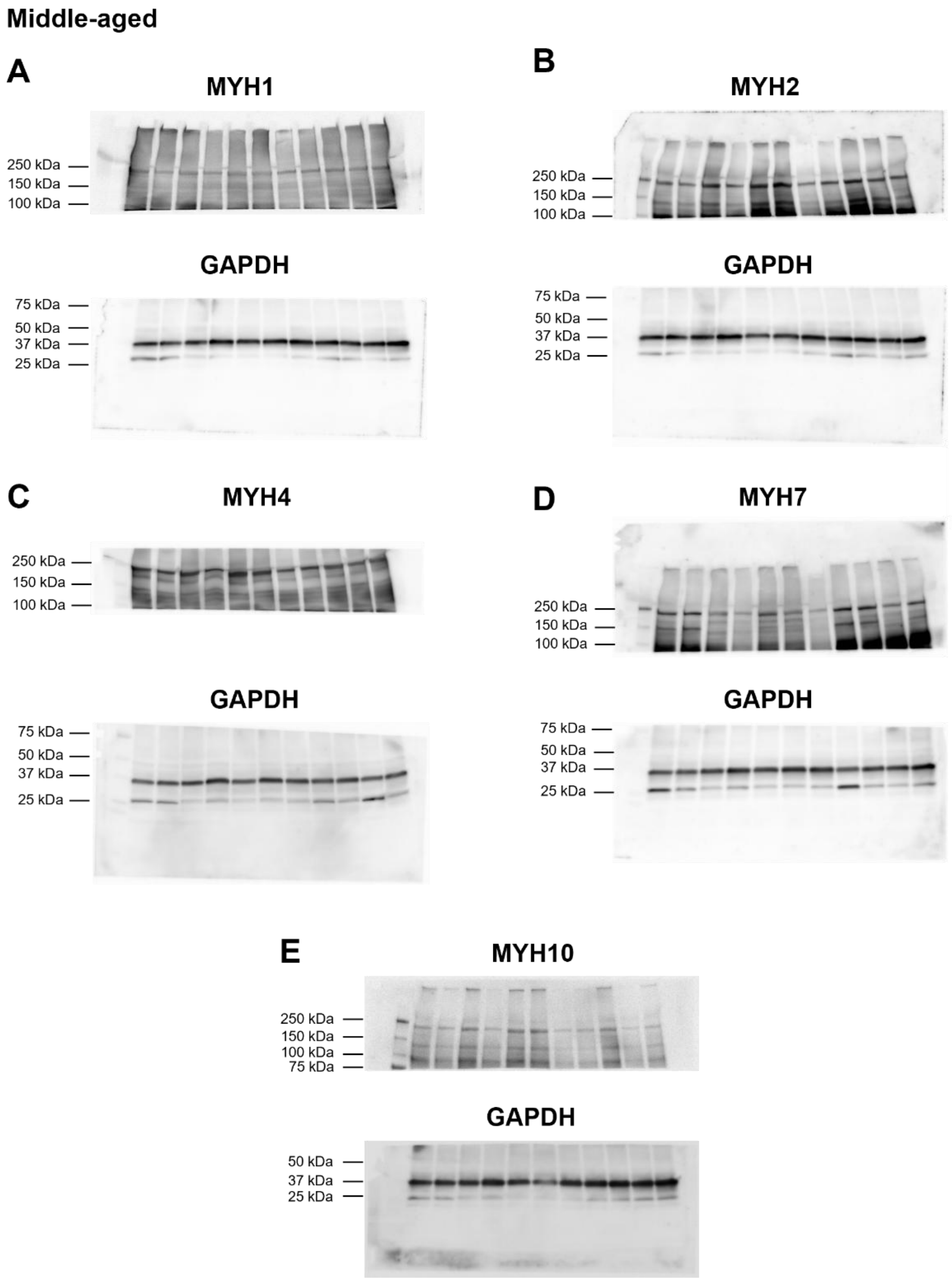


**Supplementary Figure 8. Uncropped Western blot images corresponding to Figure 6.** Uncropped Western blots of MYH1 (**A**), MYH2 (**B**), MYH4 (**C**), MYH7 (**D**), and MYH10 (**E**) from gastrocnemius muscles of 12-month-old (middle-aged) wild-type (WT) and *App*-KI mice, with the corresponding GAPDH blots. Lane order: molecular weight marker, WT-1, *App*-KI-1, WT-2, *App*-KI-2, WT-3, WT-4, *App*-KI-3, *App*-KI-4, WT-5, *App*-KI-5, WT-6. Mouse identifiers are consistent across panels. MYH, myosin heavy chain; GAPDH, glyceraldehyde-3-phosphate dehydrogenase.

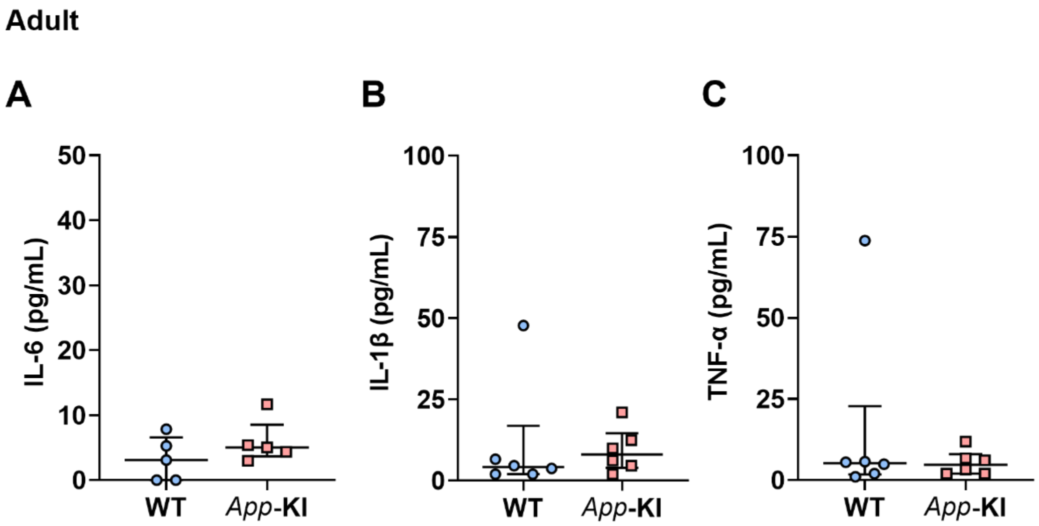


**Supplementary Figure 9. Serum cytokine levels in adult *App*-KI mice.** Serum concentrations of (A) IL-6, (B) IL-1β, and (C) TNF-α in 7–8-month-old (adult) wild-type (WT) and *App*-KI mice. Data are shown as the median with interquartile range; $n$ = 5 per group for (A) and $n$ = 6 per group for (B) and (C). Data were analyzed using the Mann–Whitney $U$ test. IL-6, interleukin-6; IL-1β, interleukin-1β; TNF-α, tumor necrosis factor-α.
